# Social and neuroendocrine phenotypes reprogrammed by endocrine-disrupting chemicals can be mitigated by *Limosilactobacillus reuteri* modulation of the gut microbiome–thyroid–oxytocin axis

**DOI:** 10.64898/2026.07.18.739173

**Authors:** Elena V. Kozlova, Maximillian E. Denys, Anthony E. Bishay, Elyza A. Do, Rui Lui, Crystal N. Luna, Artha Lam, Varadh Piamthai, Ansel Hsiao, Margarita Curras-Collazo

## Abstract

**Introduction:** Environmental factors are increasingly implicated in the etiology of autism spectrum disorder (ASD). Polybrominated diphenyl ethers (PBDEs) are anthropogenic toxicants added as flame retardants to consumer products that have become ubiquitous environmental contaminants and disrupt thyroid hormone (TH) and neuroendocrine systems. We have previously shown that developmental PBDE exposure produces ASD-like traits with involvement of oxytocin (OXT)-thyroid hormone signaling. *Limosilactobacillus reuteri* (LR), a widely used probiotic bacterium, has been shown to improve social functioning and increase TH and OXT levels in murine models. Therefore, we tested the hypothesis that LR supplementation (LR) prevents PBDE-induced deficits in socioemotional behavior with concomitant modulation of TH signaling genes on hypothalamic OXT neurons.

**Methods:** C57BL/6N mouse offspring were exposed to a commercial penta-mixture of PBDE congeners, DE-71, at an environmentally realistic concentration, 0.1 mg/kg/d (DE-71), or to corn oil vehicle (VEH/CON) via their mothers during gestation and lactation. Offspring received supplementation with LR ATCC PTA 6475 (10^7^-10^8^ CFU/mL, po) indirectly via the dam or continuation directly through adulthood. Unsupplemented controls were given saline.

**Results:** Fecal microbiome analysis in offspring confirmed colonization of LR at postnatal day (P) 40 and depletion by P104. LR treatment increased plasma total thyroxine in DE-71 and plasma OXT in VEH/CON dams. In DE-71 offspring of both sexes, LR normalized deficient scores on social novelty preference and emotional recognition in adult females and males and deficient long-term social recognition memory (SRM) in adult DE-71 females; DE-71 males were normal. Reduced olfactory dishabituation between two social odors may partly explain the compromised socioemotional behavior produced by DE-71 in an LR-dependent manner. Multiplex RNA *in situ* hybridization performed on immunoreactive OXT-ergic neurons in the paraventricular hypothalamic nucleus (PVH) revealed significant upregulation of TH transporter monocarboxylate transporter 8 (*Mct8*) and downregulation of iodothyronine deiodinase 3 (*Dio3*) in DE-71 relative to VEH/CON females. This toxicant-induced reprogramming was prevented by probiotic treatment. DE-71 males expressed reduction in *Mct8* and *Dio3* transcripts on OXT-ergic neurons with minimal LR protection. In the female supraoptic nucleus (SON), *Mct8* and *Dio3* were downregulated by DE-71 and normalized in DE-71+LR; there were no group effects on transcript levels in male SON. Results of fecal 16S rRNA sequencing indicated reduced α-diversity and altered β-diversity in the gut bacterial community of female but not male DE-71 exposed offspring; most changes were correctable by LR. Alterations in taxa-level abundance caused by DE-71 and reversed by LR were observed in both sexes. These involved *Bifidobacterium, Coprococcus, Desulfovibrio, Oscillospira*, and Peptococcaceae in females and Desulfovibrionaceae, *Rikenella*, and *Turicibacter* in males. Exposed dams showed no detriment in α- and β-diversity while showing reduced abundance of several Firmicutes and Proteobacteria taxa that could be rescued by LR. The relative abundance of *Lactobacillus* was upregulated in DE-71 males and DE-71+LR males and dams.

**Conclusions:** These results indicate that developmental probiotic supplementation effectively mitigated organohalogen-induced ASD-like deficits in socioemotional behavior and partially corrected dysbiosis of gut bacterial communities in exposed offspring of both sexes. Concomitantly, PBDEs altered the expression of TH regulatory genes *Mct8* and *Dio3* in PVH OXT neurons in a sex-dependent manner, suggesting that TH regulation of OXT neuroendocrine cells may modulate the emergence of toxicant-induced ASD-relevant behavior. While LR reinstated normal behavioral outcomes in PBDE-exposed offspring of both sexes, coincident normalization of hypothalamic TH signaling transcripts occurred more broadly in females, indicating the existence of unique parallel processes influencing the preventive effects of LR on ASD-relevant behavioral deficits in both sexes.

## Introduction

The prevalence of the neurodevelopmental disorder (NDD) Autism Spectrum Disorder (ASD) has increased substantially over the last several decades, with current estimates indicating 1 in 31 US children are affected with a 3.4:1 male: female bias (Shaw et al., 2025). ASD is diagnosed clinically solely based on deficits across two core behavioral domains: social communication and interaction, and restricted, repetitive patterns of behavior, interests, or activities (American Psychiatric Association, 2013; Lord et al., 2018). In addition to core domains, the majority of autistic individuals have co-morbidities, including intellectual disability, olfactory impairments, and gastrointestinal disturbances that are associated with microbial dysbiosis (Khachadourian et al., 2023; Larsson et al., 2017; Vuong & Hsiao, 2017). There are currently no FDA-approved pharmacological treatments to cure the core behavioral manifestations of ASD because mechanisms underlying the core symptom domains are elusive (Hirota & King, 2023), highlighting an urgent need to identify novel therapeutic approaches (Baribeau & Anagnostou, 2022). Although genetic underpinnings as well as improvements in diagnostic sensitivity and greater societal awareness contribute to ASD prevalence, they do not fully explain its continued rise (Hertz-Picciotto & Delwiche, 2009). Consequently, this has increased interest in identifying modifiable environmental risk factors, particularly prenatal toxicant exposures and maternal health status, that may interact with genetic susceptibility during neurodevelopment. Among these, polybrominated diphenyl ethers (PBDEs) have emerged as chemicals of concern because of their widespread human exposure and developmental neurotoxicity.

PBDEs are persistent indoor toxicants widely used in consumer products that readily leach into indoor environments, resulting in chronic human exposure (Herbstman et al., 2010). PBDEs cross the placenta, accumulate in blood and breast milk, raising concerns regarding their developmental neurotoxicity during early developmental windows of biological plasticity (Costa & Giordano, 2007; Scoville et al., 2019). Consistent with the developmental origins of health and disease hypothesis (DOHaD) such exposures may lead to adverse outcomes later in life (D. J. Barker, 1995; Heindel & Vandenberg, 2015). Although PBDE production has declined following regulatory restrictions, these compounds remain environmentally persistent and are predicted to continue circulating globally until at least 2050 (Abbasi et al., 2019).

Human and animal studies have identified endocrine disruption of thyroid hormones (THs) as a major health consequence associated with PBDE exposure, with additional evidence of reproductive and developmental toxicity and neurotoxicity (Costa & Giordano, 2007). Epidemiological studies have found that developmental PBDE exposure is positively associated with reduced social competence as well as poor executive function, lower IQ (Braun et al., 2014; Ding et al., 2015; Gascon et al., 2011; Gibson et al., 2018; Messer, 2010). Likewise, murine models using brominated flame retardants show disrupted social cognition, although findings are inconsistent (Gillera et al., 2020; B. Kim et al., 2015; Witchey et al., 2020; Woods et al., 2012). We recently reported that developmental exposure to the human-relevant PBDE mixture DE-71, at environmentally realistic concentrations, induces ASD-like traits in male and female mouse offspring, accompanied by altered prosocial gene expression and reduced oxytocin (OXT) immunoreactivity in the hypothalamic paraventricular nucleus, a key brain region regulating social behavior (Bielsky et al., 2004; Kozlova, Denys, et al., 2025; Kozlova et al., 2022; Kozlova, Gonzalez, et al., 2025). Collectively, these findings suggest that PBDE exposure disrupts neural circuits governing social behavior, but the mechanisms linking toxicant exposure to ASD-relevant phenotypes remain poorly understood.

One emerging mechanism involves disruption of the gut microbiome. PBDE exposure is associated with changes in the microbiome (Iszatt et al., 2019; Laue et al., 2019; Li et al., 2018; Qiu et al., 2022; Scoville et al., 2019) and early life disruption of the microbiome by xenobiotics has been linked to behavioral disorders and the onset of human disease later in life (Clarke et al., 2019; Tamburini et al., 2016). Recent paradigm-shifting studies in animal ASD models have shown that certain ‘psychobiotic’ gut bacteria can modulate central nervous system-driven behaviors, including the elimination of social behavior deficits relevant to ASD (Buffington et al., 2016; Dinan et al., 2013; Hsiao et al., 2013; Morais et al., 2020; Sgritta et al., 2019). Among these, lactic acid bacteria species *Limosilactobacillus reuteri* (ATCC PTA 6475) has emerged as a particularly promising candidate because of its ability to influence OXT signaling through the gut–brain axis.

Notably, OXT is altered in autistic individuals and exogenous application alone or with the OXT- and TH-promoting probiotic *L. reuteri* has been used in the treatment of ASD with mixed results (Audunsdottir et al., 2024; Huang et al., 2021; Mazzone et al., 2024). LR (ATCC PTA 6475) can also influence OXT signaling through the gut-brain axis, especially in regulating social behavior (Cuesta-Marti et al., 2023; Danhof et al., 2023; Erdman, 2014). OXT neurons in the PVH are likely critical for social and emotional recognition ability, via their projections to various oxytocin receptor-rich forebrain regions (Ferretti et al., 2019; Thirtamara Rajamani et al., 2024). Postnatal administration of LR ATCC PTA 6475 can increase OXT levels in the PVH in various mouse models of ASD, and rescue ASD-relevant behavior (Buffington et al., 2016; Dinan et al., 2013; Hsiao et al., 2013; Morais et al., 2020; Sgritta et al., 2019). Previous reports also demonstrate the TH promoting and immune bolstering bioactivity of LR 6475 (Lin et al., 2008; Varian et al., 2014). Therefore, PBDE-induced ASD-like behavior may be due, in part, to disruption of the gut microbiome and downstream reduced OXT signaling via documented disruption of thyroid hormone functions. However, no studies to date have examined the participation of and potential manipulation of the gut microbiome–thyroid–oxytocin axis to counteract toxicant-induced ASD-like phenotypes. We therefore hypothesized that developmental PBDE exposure disrupts the gut microbiome–thyroid–oxytocin axis, resulting in ASD-relevant behavioral abnormalities, and that postnatal supplementation with *L. reuteri* would mitigate these effects.

Here, we investigated the long-term effects of maternal DE-71 exposure and *L. reuteri* supplementation on ASD-relevant parameters in exposed C57BL/6 mice offspring. We assessed performance on social recognition memory, emotional recognition, and social olfactory discrimination tasks to characterize ASD-relevant phenotypes in the presence and absence of LR treatment. Using multiplex RNA in situ hybridization, we also examined TH regulatory gene markers expressed on immunofluorescence-labeled hypothalamic OXT-producing neurons to explore potential pathways underlying toxicant endocrine disruption and probiotic-mediated protection. Next-gen sequencing was used to determine the effects of DE-71 exposure and LR probiotic treatment on maternal and offspring gut microbial communities. Together, these investigations were designed to provide new insights into the neuro/endocrine basis of susceptibility to neurological reprogramming by developmental toxicant exposure and to elucidate the potential benefits of LR probiotic treatment.

## Methods

### Animal Care and Maintenance

C57BL/6N mice (Charles River Laboratories; Raleigh, NC, USA were group-housed 2-4 per cage on corn cob bedding. Mice were maintained in a specific pathogen-free vivarium on a 12:12 h light:dark cycle at 20.6-23.9°C and relative humidity of 20-70%. Mice were provided Laboratory Rodent Diet (5001; LabDiet, Quakertown, PA, USA) and tap water *ad libitum*. Procedures on animals were performed in compliance with the National Institutes of Health *Guide for the Care and Use of Laboratory Animals* and approved by the University of California, Riverside, Institutional Animal Care and Use Committee under AUP# 5 and 20210031.

### Developmental exposure to DE-71

Perinatal PBDE exposure via the dam was accomplished as described previously (Denys et al., 2025). In brief, DE-71, a pentabromodiphenyl ether mixture (Lot no. 1550OI18A; CAS 32534-81-9; Great Lakes Chemical Corporation, West Lafayette, IN, USA) was dissolved in corn oil (Mazola; Summit, IL, USA) and sonicated for 30 min to yield a low dose at 0.1 mg/kg bw/day (2 µL/g bw). Controls received corn oil without DE-71 (VEH/CON).

Dams were fed corn flakes infused with DE-71 or vehicle solution daily, except on postnatal day (P) 0 and 1 for 10 weeks that included ∼4 weeks of pre-conception, plus gestation (3 weeks) and lactation (3 weeks). This dose and regimen were chosen to model human-relevant chronic, low-level exposure (Costa & Giordano, 2007; Drage et al., 2019; Han et al., 2020; Ongono et al., 2019). LC/MS analyses conducted by our team detail the resulting levels and composition of PBDE congeners in brain tissue of mice offspring exposed in this way (Kozlova et al., 2020, 2022; Kozlova, Gonzalez, et al., 2025). During the last week of the 4-week pre- conception exposure period, dams were mated with an untreated C57BL/6N male. Offspring were weaned after the lactation period at P21. Dams were examined for perinatal parameters, maternal behavior, plasma endocrine hormones and fecal microbiome composition. Female and male offspring were studied for neurobehavior and *ex vivo* analysis of plasma oxytocin, brain oxytocin receptor and TH markers using dual immunofluorescence/RNA *in situ* hybridization and fecal microbiome composition. The experimental timeline is shown in **Fig. 1A**.

**Figure 1.**
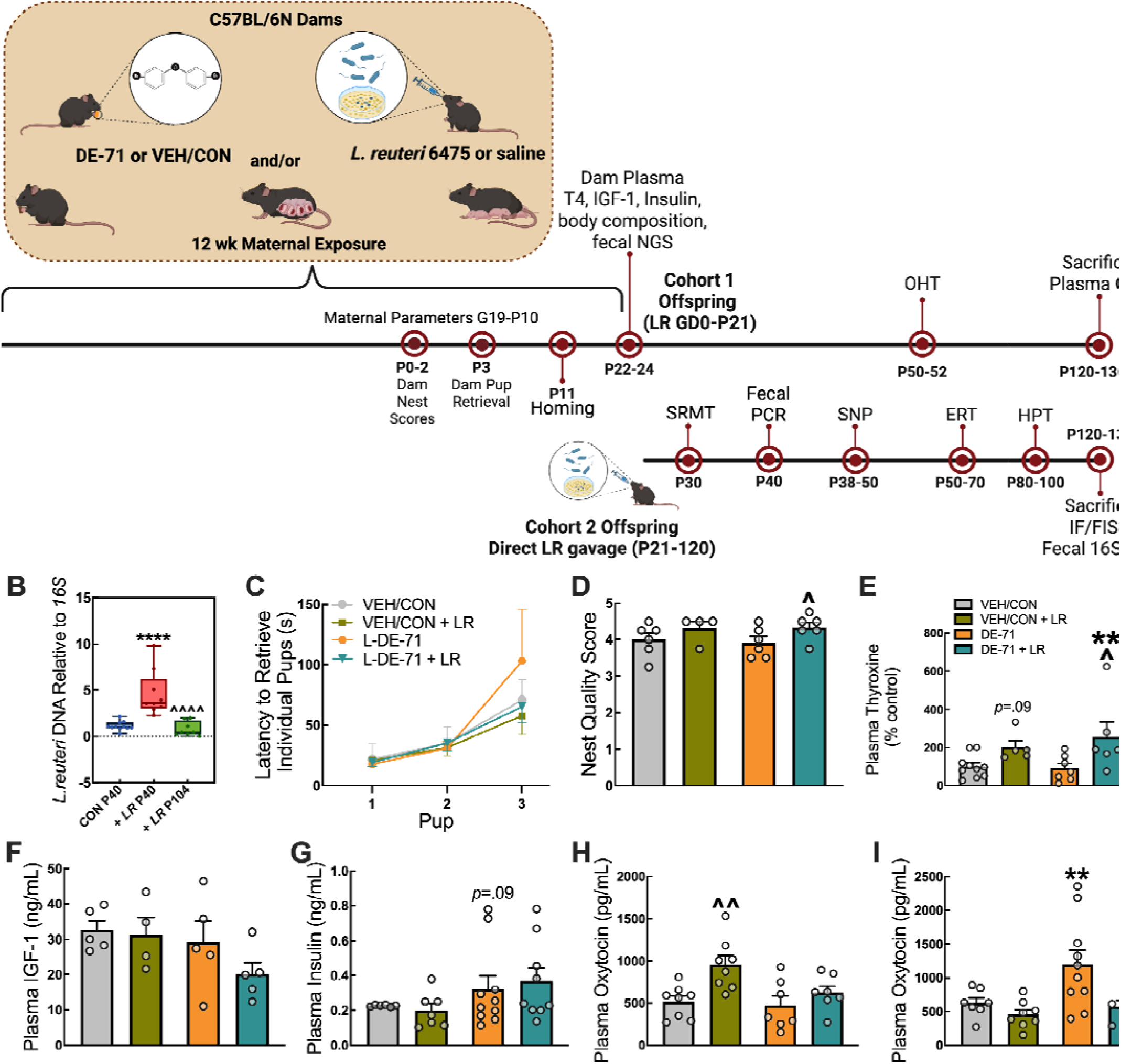
Experimental timeline of PBDE exposure and probiotic administration and parameters measured in dam mothers and their offspring. (**A**) Direct oral exposure of dams to a mixture of PBDE congeners, DE-71 (0.1 mg/kg/d), or vehicle control (VEH/CON), began ∼3 weeks before conception and continued daily through gestation and lactation until pup weaning at P21. A total of four treatment groups received DE-71 or VEH with either supplementation of *L. reuteri* ATCC PTA 6475 (∼1×10^7^-10^8^ CFU/mL/d) or saline (unsupplemented). LR treatment was provided via oral gavage daily to dams on the first night of mating until P21 (Cohort 1) or was additionally given directly to offspring from weaning until sacrifice (Cohort 2). Offspring behavior was examined at P11 (homing - maternal attachment), at P28 (SRMT) and as adults (various). Dams were assessed for plasma, body composition and fecal NGS after pup weaning. (**B**) Fecal colonization of *L. reuteri* ATCC PTA 6475, normalized to total bacterial 16S rDNA, was measured by RT-qPCR in offspring at P40 after receiving 18 doses of LR or at P104 compared to control (CON). (**C**) Latency for dams to retrieve each displaced pup back to the nest at P3. (**D**) Dam nest quality scores during P0-2. (**E**) Dam plasma total T4 at P24. (**F**) Dam plasma IGF-1 at P24. (**G**) Dam plasma insulin at P24. (**H,I**) Offspring plasma OXT of female (H) and male offspring (I) at sacrifice. Whisker plots in Panel B represent median and interquartile range representing minimum and maximum values. Bar plots represent mean +/- sem values. *statistical difference vs VEH/CON P40 (B) or VEH/CON (E,I), \*\**p*<0.01, \*\*\*\**p*<0.0001. ^statistical difference vs LR P40 (B) or corresponding unsupplemented group (D,E,H,I), ^*p*<0.05, ^^*p*<0.01, ^^^^*p*<0.0001. *n*, 4-10/group, IF: immunofluorescence; T4: total thyroxine OXT: oxytocin; P: postnatal day; GD: postnatal day; IF/FISH: dual immunofluorescence/RNA *in situ* hybridization; NGS: next-gen sequencing; SNP: social novelty preference test; SRMT: social recognition memory test; OHT: olfactory habituation/dishabituation test; ERT: emotional recognition test; CFU: colony-forming units; LR: *Limosilactobacillus reuteri*

### Monospecies Probiotic Culture and Treatment

*Limosilactobacillus reuteri* (LR, formerly *Lactobacillus reuteri* (Zheng et al., 2020)) ATCC PTA 6475 (gift of BioGaia, Stockholm, Sweden) was cultured anaerobically with deoxygenated De Man, Rogosa, and Sharpe (MRS) media as described (Denys et al., 2025). Dams were orally gavaged with LR or saline daily beginning on the first night of mating until weaning of pups at P21. Cohort 1 mice offspring received LR via mother only until weaning at P21, while Cohort 2 offspring continued to receive LR or saline via oral gavage daily until sacrifice.

### Maternal Parameters

#### Dam Food and Water Intake

Dam food and water intake, gestational weight gain and litter size and sex ratio were monitored (**Supplementary Table 1**). Food and water were weighed daily at the time of dosing. The pups were counted and sexed on the day of birth and secondary sex ratio calculated as number of males over total pups in litter.

#### Nest scoring

The nests of single-housed dams built from pressed cotton squares were evaluated when litters were P 0-1 using a modified scoring system based on the height and closure of the walls surrounding the nest cavity (Hess et al., 2008). Pictures of the nests were taken between 9:00 and 16:00 and scores were assigned by at least two observers. Scores were compared according to the following rubric: nest contained a center (1) plus a 50% border (2) or 75% border (3) or 100% border (4). Nest scores were boosted by 0.5 if the nest walls were elevated. A score of 5 was given if the nest had higher than 1 inch walls and resembled a dome. Nests with pups strewn about outside the nest received a score of 0. Bland-Altman plots indicate agreement and negligible skewing in scores provided by separate observers (**Supplementary Fig. 1**).

#### Pup Retrieval Test

Maternal behavior was assessed using a pup retrieval test on postnatal day (P)3-4. All offspring were separated from the dam’s nest for 20 min into a separate cage and kept on a heating pad to maintain body temperature. Three randomly selected pups were returned to the dam’s cage and placed in the corners. Latency to investigate the first pup and duration taken to return each of 3 pups to the home nest was scored by analyzing digital recordings of the test (Wu et al., 2009). The total test duration was limited to 25 min at which time the litter was returned to the homecage.

#### Neurobehavioral Testing Paradigms

Mice were acclimated to the testing room and apparatus at least 30 min prior to testing. Testing apparati were cleaned with soap and water in between individual mouse trials to remove debris and odors. Unless stated otherwise, mouse behavior was scored using automated video-tracking software (Ethovision XT 17, Noldus Information Technologies, The Netherlands) or manual event-logging software (BORIS (Friard & Gamba, 2016)), by trained observers blinded to treatment. Mice were tested between 10 am and 4 pm during the light phase under bright light conditions, unless otherwise stated.

#### Maternal Attachment Test (Homing)

The homing test is based on a neonatal pups tendency to remain in close contact with the dam and siblings, which requires cognitive ability for differentiating between own vs other home cage odors as well as requires olfactory and motor abilities to navigate toward the mother’s odor (Bignami, 1996; Scattoni et al., 2008). On P11, the floor (26 cm length × 16 cm width × 12 cm height) of a clean Plexiglass mouse cage was subdivided into three areas by wire-mesh dividers. One area was uniformly covered with soiled corn-cobb bedding and nestle material from the home cage. Soiled bedding from a DOB-matched litter of a stranger dam cage were placed on the opposing one third of the cage. The middle third of the cage was covered with clean bedding. Pups were placed individually in the middle section for 1 min, the dividers were then removed and the pups were allowed to explore the arena for 2 min. The total time the center point of the pups spent was scored by Ethovision.

#### Social Recognition Memory Test (SRMT)

Social recognition memory test measures long-term recognition memory and was performed as described previously (Kozlova et al., 2022). Briefly, adult test mice were placed in a standard mouse cage for 45 min under dim and quiet conditions. After acclimation, the adult test mouse was exposed to a single juvenile sex-matched conspecific (P11-P13) twice for (3 min) at an interval of 24 h and time taken to investigate the juvenile during day 1 (Trial 1) and day 2 (Trial 2) was assessed. An interaction of 3 min was digitally recorded during each Trial. Stimulus mice were not used more than 3 times/day. Investigation times were later quantified with event-logging BORIS software used for behavioral observation research. Test mouse behaviors that qualified as social investigation were inspecting any part of the body including licking, pawing, grooming, and even sniffing the tail while being at least 1 cm away from the stimulus mouse. Aggressive and chasing encounters were disqualified.

#### Social Novelty Preference (SNP)

Formation of short-term social recognition memory was tested using a social novelty preference (SNP) test conducted using modifications from described protocols (Moy et al., 2004). Test mice were habituated to the testing room and empty large test cage (46 cm L × 24 cm W × 16.5 cm H) for 30 min. Mice were exposed to 2 wire interaction corrals (11 cm height × 9.5 cm inner diameter) placed at each side of the cage for 30 min. During a 5-min training trial, a sex- and age-matched stimulus mouse was placed into one corral while the empty corral was removed. After a 30 min retention period, the test mouse was introduced to the same stimulus mouse (now familiar mouse) and a novel stimulus mouse for 5 min. Prior to testing days, stimulus mice were trained to stay in corrals. Social recognition memory (SRM) was represented as significantly greater time spent investigating novel vs familiar stimulus mouse as percent of total investigation time.

#### Emotional Recognition Test (ERT)

An emotional discrimination test (ERT) was administered to determine if PBDE mice are able to discriminate unfamiliar conspecifics based on their negatively valenced emotional state as described (Ferretti et al., 2019; Scheggia et al., 2020). Mice were habituated to the testing cages for 10 min. for three days leading up to the first experiment. The test mice were habituated inside a polycarbonate cage (27 x 16 x 12 cm) with a polyvinyl separator and two cylindrical metal wire cups. Sex and aged matched ‘demonstrator’ mice were habituated inside the testing cages, under the wire cups three times for 10 minutes. Demonstrators were re-used for maximum two/three times, with at least 5 days between each test. Stress demonstrators were generated by placing mouse in a restrainer for 15 min immediately prior to the onset of the testing session. Each test was digitally recorded and scored with Ethovision software.

#### Olfactory Habituation/Dishabituation Test

The ability of mice to detect and differentiate social and non-social odorants was examined using the olfactory habituation/dishabituation test (OHT) (Silverman et al., 2010). During OHT test mice are presented with a cotton-tipped applicator infiltrated with non-social and social odorants in 2-min triplicate trials in the following order: water, almond, banana, social odor 1, social odor 2. Mice were acclimated for 45 min to an empty standard mouse cage with an untreated cotton-tipped applicator inserted through the water bottle hole on the metal lid. Non-social odors were prepared from water and almond and banana extracts (1:100; McCormick) immediately before testing. Two different social odors were obtained the morning of test day by swiping applicators across the bottom of two occupied mouse cages containing soiled bedding from unrelated sex-matched conspecifics. Soiled cages housed 3–4 mice and bedding was at least 3d old. Time spent sniffing the infiltrated applicator was recorded with a stopwatch. Parameters measured were habituation, defined as a decrement in olfactory investigation of the same odor after repeated presentations and dishabituation, defined as a reinstatement of olfactory investigation upon presentation of a new odorant.

#### Hot Plate Test

We measured an animal’s tolerance to a thermal stimulus and pain threshold using a hot plate test using a modification of a previously published protocol (Woolfe & Macdonald, 1944). Briefly, a cast iron steel pan (1.9 cm thick) was placed in a water bath over a hot plate. The surface temperature was maintained at 53 ± 1.0 °C as measured by an infrared thermometer (General IRT-207). The animals were monitored and the latency to exhibit nociceptive responses (i.e. hindpaw withdrawal or licking, jumping) was recorded (Minett et al., 2011). A predetermined cut-off time after which the test was stopped was set at 30 s to prevent tissue damage.

#### Dual Multiplex Single-molecule RNA Fluorescence *in situ* Hybridization (smFISH) and Immunofluorescence (IF)

Mice were sacrificed using CO_2_ inhalation and perfused transcardially (exsanguinated) with 0.01 M phosphate-buffered saline (PBS) followed by 4% paraformaldehyde (PFA) in 0.1 M phosphate buffer (PB). After 24 h post-fix brains were cryoprotected in successive incubations of 10, 15 and 30% sucrose in PBS over 3 d. Solutions were treated with 1% diethyl pyrocarbonate (DEPC). Brains were stored at -80 °C until cryosectioned on a sliding cryostat (Leica Microsystems, Germany) at 20 μm thickness. Sections were mounted on adhesive microscope slides and fluorescent labeling of mRNA transcripts (RNAscope) was performed using the Multiplex Fluorescent Reagent Kit V2 Assay (323100; Advanced Cell Diagnostics, Hayward, CA, USA) according to the manufacturer’s instructions. In brief, sections were air-dried at RT for 20 min before post-fixation in 4% PFA (30 min), baking at 60 °C (60 min), dehydration through graded ethanol solutions (50, 70, 100%), treatment with H_2_O_2_ (10 min), target retrieval at 94 °C (5 min) and with protease III treatment at 40 °C (30 min). Sections were subsequently hybridized with probes for 2 h at 40 °C in a humidified oven (HybEZ II, ACD-Bio, Newark, CA, USA): Mm-Dio3-C1 (*Mus musculus* deiodinase iodothyronine type III, cat. no. 561641), Mm-Slc16a2-C2 (Mct8) (*Mus musculus* solute carrier family 16 member 2, cat. no. 545291-C2),), Mm-Oxtr-C1 (*Mus musculus* Oxytocin receptor, cat. no. 412171). Positive and negative controls were used: Probe-Mm-(Polr2a-C1, PPIB-C2, Ubc-C3) (cat. no. 320881) and Probe-Mm-DapB-C1/2 (cat. no. 320871), respectively. Sections on slides were then incubated with Multiplex FL v.2 Amp1 (30 min), Amp2 (30 min), and Amp3 (15 min). The fluorescent signal was developed by consecutive incubation with Opal dyes from Akoya Biosciences: Opal 690 ([*Dio3* (C1 channel)], FP1497001KT) or Opal 620 ([*Mct8* (C2 Channel, *Oxtr* (C1 channel)], FP1495001KT).

Sections were then incubated with mouse monoclonal anti-oxytocin neurophysin primary antibody (PS38, 1:100; gift of the late Dr. Harold Gainer, NIH, Bethesda, MD USA) for 24 h at 4°C. The PS38 antibody is highly specific for the oxytocin-associated carrier protein Neurophysin I (Ben-Barak et al., 1985), (Whitnall et al., 1985). Following washes in PBS sections were incubated with goat anti-mouse Alexa Fluor 488 secondary antibody (1:1000, Life Technologies, USA) at RT for 1.5 h. Slides were coverslipped with antifade mounting media (Vectashield Plus, Vector Laboratories, Burlingame, CA USA) and cover glass (Precision, Cat # CG15KH1, Thorlabs, Newton, NJ, USA). Immunofluorescence images were acquired on an Zeiss Axio Imager M2 fluorescence upright microscope (Carl Zeiss, Germany) or Evident FLUOVIEW FV4000 Confocal LSM.

#### Automated Quantification of smFISH and IF using Artificial Intelligence (AI) Powered Analysis Pipeline

mRNA puncta were counted using 4-8 unilateral microscope fields per animal and at least three independent animals. Channels within each image were visually inspected for background noise, artifacts, and signal integrity. Images that failed to meet quality standards (tissue containing tears, folds or staining artifacts) were excluded from analysis. Images were analyzed for sub-cellular mRNA puncta on DAPI or /OXT-stained cells using the HALO-AI digital image analysis software (v3.6, Indica Labs, USA). HALO-AI is a trainable program that uses deep-learning neural network algorithms to accurately segment individual cells that are not always of uniform size. ROIs drawn over each anatomical field from each experimental group were chosen for training with HALO-AI deep learning nuclear segmentation using the Nuclei Seg V2 (DAPI) and Membrane Seg (*Oxt*) classifiers. Training was performed at a resolution of 0.32 um/px until a stable cross entropy value was observed. Classifiers were subsequently fine-tuned within the Nuclear Detection parameters to standardize identification across all treatment groups. For counts of DAPI-specific puncta, a constant 2 um was used within the cell expansion feature to allow puncta to be associated per cell. This was not necessary for OXT-positive neurons. The quality of the generated nuclear segmentation was evaluated manually by 3 experimenters. The High Plex FL (v4.1.3) module in HALO was then used for subcellular detection of mRNA transcripts.

#### Quantification of smFISH using QuPath

Images were acquired of 4–8 microscope fields from bilateral CA2 field of the hippocampus from each test mouse using a 20x/0.5 NA objective (EC Plan-NEOFLAUR, Zeiss) on a Zeiss Axio Imager M2 upright microscope (Carl Zeiss, Germany) equipped with a Hamamatsu ORCA-Flash 4.0 V3 digital camera. Images were analyzed using open source software (QuPath v0.2.3 (Bankhead et al., 2017)). Cell detection was used to quantify the number of DAPI-stained nuclei followed by sub-cellular detection to quantify mRNA transcripts.

#### Plasma Thyroid Enzyme-linked Immunosorbent Assays (ELISA)

Plasma was isolated from blood collected from dams via cardiac puncture at sacrifice (*ad libitum* fed state) and total thyroxine (T4) was quantified using a commercial kit following manufacturer instructions (K050-H1, Arbor Assays). Sensitivity and dynamic range of the assay were 0.3 ng/mL, 0.63-20 ng/mL, respectively. Of note, the thyroxine antibody in the kit had an 89% reactivity with rT3.

#### Insulin-like Growth Factor ELISA

Plasma was collected from post-lactating dams at sacrifice two days following pup weaning (ad libitum fed state) and assayed for Insulin-like Growth Factor (IGF-1), a peptide involved in CNS growth and development that also improves insulin sensitivity (Friedrich et al., 2012), using a commercial kit according to the manufacturer’s instructions (Cat.#EMIGF1, ThermoFisher Scientific, Waltham, MA). Sensitivity and dynamic range of the kit was 4 pg/mL and 2.74-2,000 pg/mL, respectively. The assay has no cross-reactivity with 61 cytokines and peptides tested including leptin.

#### Insulin ELISA

Plasma collected from post-lactating dams at sacrifice (ad libitum fed state) was analyzed for resting plasma insulin concentration using an ALPCO Ultrasensitive Insulin ELISA kit (Cat.# 80-INSMSU-E01, Salem, NH) with a sensitivity of 0.019 ng/mL in a standard range of 0.025-1.25 ng/mL.

#### Oxytocin ELISA

Blood was collected by cardiac puncture and the plasma separated at 2000_×_g centrifugation for 20 min at 4_. Plasma levels of oxytocin were quantified using commercially available multi-species ELISA kit from Arbor Assays (K048-H1; Ann Arbor, MI USA) following the manufacturer’s instructions. The kit had a sensitivity of 1.7 pg/sample in a dynamic range of 16.38–10,000 pg/mL. Plasma oxytocin was quantified by interpolating absorbance values using a 4-parameter-logarithmic standard curve (MyAssays).

#### Fecal DNA Isolation and Bacterial Quantification with qPCR

Fecal pellets were obtained from Cohort 2 offspring at P40 and disrupted mechanically to yield lysates from which DNA was isolated. Purity and quantity of DNA was assessed photometrically using 260/280 nm and 260/230 nm ratios. Oligonucleotide PCR primers for *L. reuteri* and the universally conserved bacterial 16S ribosomal RNA gene, obtained from Integrated DNA Technologies, Inc. (Coralville, IA, USA), were 97.5-100.7% efficient in detecting DNA extracted from LR cultures. The PCR protocol used has been published elsewhere (Denys et al., 2025). Fold expression for LR was measured relative to the universal bacteria reference gene 16S, and differential gene expression was compared to dam baseline levels or saline-supplemented controls using the Pfaffl method (Pfaffl, 2001).

#### 16S Library Preparation and Next-Gen Sequencing

Amplicon sequencing of the V4 variable region of the 16S ribosomal gene was performed as previously described (Alavi et al., 2020; Denys et al., 2025). Paired-end 150 nt reads were assembled, demultiplexed, and analyzed using QIIME 2 2023.9 software packages (Caporaso et al., 2010). Taxonomy was classified based on Greengenes (gg) 13 8 99% operational taxonomic units (OTUs). Each sample was rarefied to 1000 as the sampling depth for alpha and beta diversity analysis.

#### Statistical Analyses

Power analyses were performed to establish sample size (GPower 3.1 (Faul et al., 2007), Universitat Dufferldorf). All statistical analyses were conducted using GraphPad Prism (GraphPad Software v9.5.1). Bars and error bars in graphs represent mean±s.e.m unless indicated otherwise. Statistical analysis of main effects was accomplished using Student’s t-test, one-way, two-way or mixed model analysis of variance (ANOVA) with or without a repeated measures (RM) design or mixed effects model. Within group comparisons were performed using paired Student’s t test. Non-parametric statistical tests (i.e., Mann-Whitney, Kruskal Wallis) were used when normality assumptions were not met as measured using the Shapiro–Wilk test. If equal variance assumptions were not met using F-tests, a Welch’s t-test or Brown–Forsythe or Welch’s ANOVA were used. For two-way ANOVA, Holm Sidak’s or Fisher’s least significant difference *post hoc* tests were used. QIIME 2 2023.9 (Caparoso at al., 2010) and R (v4.0.4 Boston, MA, USA) was used for analysis of 16S rRNA sequencing data. Group comparisons of microbial community structure were performed using permutational multivariate analysis of variance (PERMANOVA), Krusskal-Wallis or Mann-Whitney ests. Differences were considered statistically significant at *p*<0.05.

## Results

### LR transfer to offspring, maternal physiology, and behavior

Fecal pellets from Cohort 2 offspring were subjected to qPCR analysis at P40 and P104. qPCR analysis of LR DNA relative to general 16S bacterial DNA showed increased expression of LR in supplemented offspring compared to saline controls (CON) (*p*<0.0001) and normalized at P104 (vs P40, *p*<0.0001) (**Fig. 1B**). Similar results have been previously shown for Cohort 1 in a report on their developmental benchmarks (Denys et al., 2025).

Food and water intake and weight gain were measured in dams throughout gestation and lactation. No effects of DE-71 exposure or LR treatment were observed. The secondary sex ratio, or number of males:females at birth, was not different across groups (**Supplementary Table 1**). However, a comparison of LR-supplemented and unsupplemented groups showed a significant reduction in the former in agreement with a previous report that suggested the involvement of an OXT-dependent mechanism (Erdman, 2014) (**Supplementary Fig. 2**).

Maternal behavior was investigated using a maternal attachment test and nest quality. There were no group differences in latency to retrieve individual pups after being experimentally displaced from the nest (**Fig. 1C**) or in nest scores (**Fig. 1D**). Maternal endocrinological parameters such as plasma total thyroxine (T4) was elevated only in DE-71 + LR (*p*<0.01, **Fig. 1E**). Similarly, plasma IGF-1 (**Fig. 1F**) showed no significant effect of exposure or treatment differences. Plasma insulin showed an *apparent* increase in the DE-71 group (**Fig. 1G,** *p*=0.09). There were no differences in dam lean and fat mass after pup weaning (P23) as a percentage of body weight (**Supplementary Figure 3**).

Since the prosocial behavior outcomes tested in this study rely on plasma OXT we examined exposure and treatment effects on plasma OXT in offspring at P150. In females, LR treatment significantly increased plasma OXT in VEH/CON+LR group (*p*<0.01, **Fig. 1H**). Male but not female DE-71 offspring displayed a significant elevation as compared to VEH/CON (*p*<0.01), an effect that was prevented in DE-71+LR (vs DE-71, *p*<0.05) and normalized (vs VEH/CON, ns) (**Fig. 1I**).

### LR supplementation prevents deficits in social and emotional recognition ability produced by DE-71

Next, we investigated the effect of DE-71 exposure and LR treatment on autism-relevant behaviors. For females on the SNP test, which measures short-term social memory, all groups except DE-71 showed increased preference to investigate a novel compared to a familiar mouse stimulus (novel vs familiar, *p*<0.05) (**Fig. 2A**). Similarly, DE-71 female offspring displayed abnormal behavior on SRMT, a test of long-term social recognition memory. The other female groups, including the LR supplemented DE-71 females (*p*<0.05), spent significantly less time investigating familiar mouse on Day 2, indicating sufficient memory for the previously encountered mouse on Day 1 (**Fig. 2C,** *p*<0.01-0.001). To test the effect of DE-71 exposure and LR treatment on emotional recognition, we placed test mice in a cage with demonstrator mice that exhibited either neutral or stress emotional states. VEH/CON female offspring spent significantly more time investigating stressed demonstrator as compared to neutral controls (stressed vs neutral, *p*<0.01), while DE-71 females spent equal time on both stimulus mice (**Fig. 2E**). Notably, DE-71 mice supplemented with LR showed normal emotional recognition (*p*<0.05), although LR supplemented VEH/CON females showed only an apparent difference (*p*<0.09).

**Figure 2.**
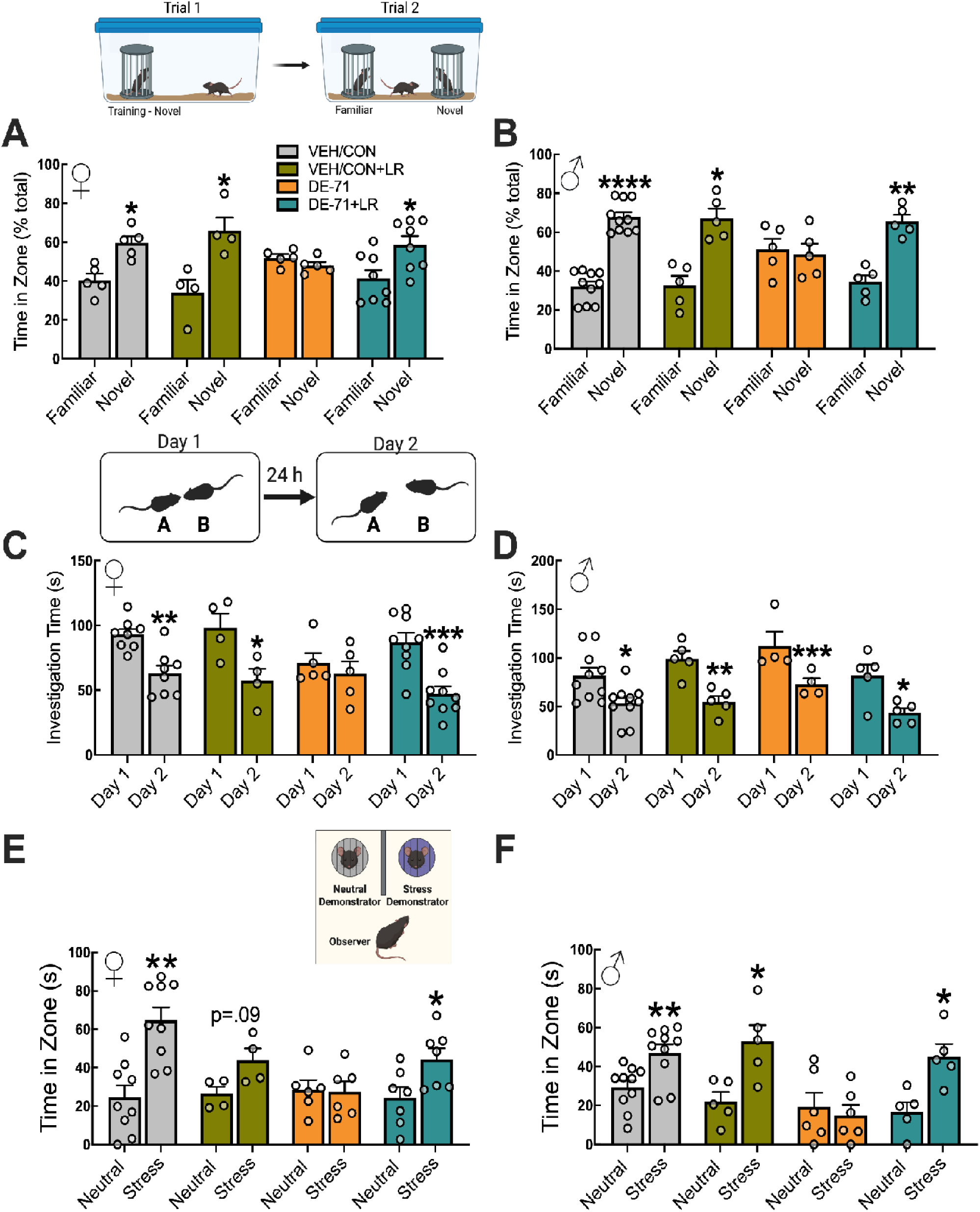
LR supplementation prevents deficits in short- and long-term social recognition ability and emotional recognition produced by DE-71. (**A,B**) Percentage of time spent in novel and familiar zones on social novelty preference (SNP) test for female (A) and male offspring (B). (**C,D**) Time spent investigating familiar stimulus mouse on days 1 and 2 of a social recognition memory test (SRMT) for females (C) and male offspring (D). (E,F) Emotional Recognition test in female (E) and male offspring (F). *statistical difference novel vs familiar stimulus (A,B) or Day 1 vs Day 2 (C,D) or stress vs neutral demonstrator (**E,F**), \**p*<0.05, \*\**p*<0.01, \*\*\**p*<0.001, \*\*\*\**p*<0.0001. *n*, 4-10/group

Similar results were obtained for male offspring on SNP test. All male groups except DE-71 spent more time investigating the novel vs familiar mouse: VEH/CON (*p*<0.0001), VEH/CON+LR (*p*<0.05) and DE-71+LR (*p*<0.01) groups (**Fig. 2B**). However, DE-71 males did not prefer novelty on SNP, indicating deficient short-term SRM. On SRMT, which measures long-term social recognition memory, all male groups, in contrast to females, showed normal behavior, i.e., less investigation of familiar stimulus mouse on Day 2 (**Fig. 2D**): VEH/CON (*p*<0.05), VEH/CON+LR (*p*<0.01), DE-71 (*p*<0.001) and DE-71+LR (*p*<0.05). On ERT male mice behaved similarly to females with all groups but DE-71 preferring stress vs neutral demonstrators (*p*<0.05-0.01). Importantly, the abnormal phenotype manifested by DE-71 was prevented by LR treatment (*p*<0.05).

In order to assess the ability of offspring to ambulate and show preference in early development we subjected them to a maternal attachment/homing test at P11. Female and male pups across all treatment groups preferred bedding with odors from their mothers relative to that from other mothers (**Supplementary Fig. 4**).

Taken together, these results suggest that developmental exposure to DE-71 produces ASD-like traits on behavioral tests representing 3 different constructs, which can be prevented by LR supplementation. One exception to this is that male DE-71 mice were not abnormal on SRMT. This is consistent with our previous report of a weakly abnormal long-term SRM phenotype in exposed male offspring (Kozlova, Gonzalez, et al., 2025).

### Perinatal exposure to DE-71 alters social odor discrimination in adult offspring; LR protection

To investigate if abnormalities in socioemotional behaviors were associated with altered odor discrimination, we subjected offspring to an olfactory habituation/dishabituation test (OHT). Graphical data is shown in **Figure 3** and statistical results are summarized in **Table 1**. We found normal habituation (*p*<0.05-0.0001) and dishabituation (*p*<0.05-0.001) to non-social odors tested in all female and male groups with one exception being habituation to water for DE-71+LR. DE-71 females showed dishabituation from last non-social odor to social odor 1 (banana 3 vs social odor 1, *p*<0.0001) but not from social odor 1 to 2 (social odor 1-3 vs social 2-1, ns). Females of all groups (*p*<0.0001) showed normal habituation to social odors (*p*<0.0001) except for DE-71 females that had incomplete habituation to social odor 2 (social odor 2-1 vs social odor 2-3, ns). Between-group comparisons for females revealed significantly reduced mean sniffing score for social odor 1 in DE-71 (*p*<0.001) that was not normalized with DE-71+LR (vs VEH/CON, *p*<0.01, **Fig. 3A**). DE-71 also displayed reduced mean sniffing score of social odor 2 as compared to VEH/CON (*p*<0.0001), an effect that was normal in DE-71+LR (vs DE-71, *p*<0.0001, **Fig. 3B**).

**Fig. 3.**
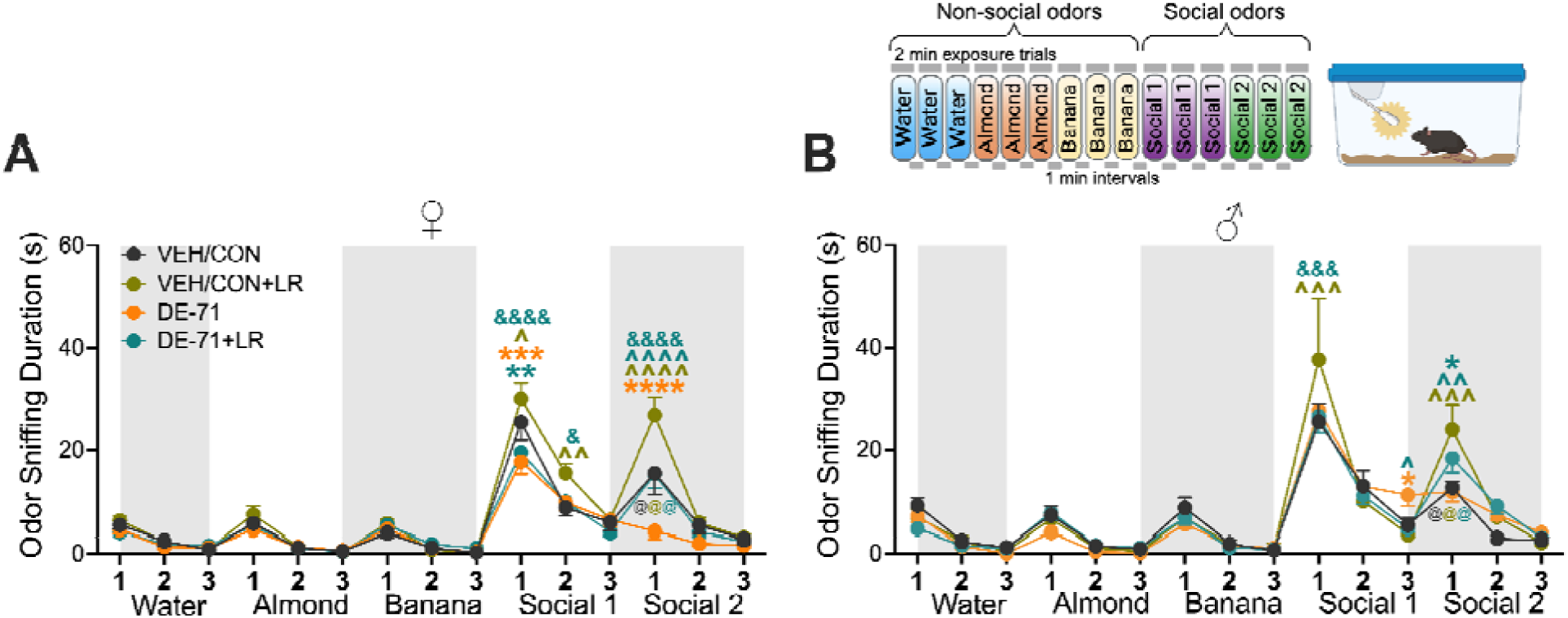
Perinatal exposure to DE-71 alters olfactory discrimination of social odors in adult offspring; protection by LR. Sniffing time of non-social and social odors measured on olfactory habituation/dishabituation test (OHT) in females (**A**) and males (**B**). *significant between-group difference vs VEH/CON, \**p*<0.05, \*\**p*<0.01, \*\*\**p*<0.001, \*\*\*\**p*<0.0001. ^significant between-group difference vs corresponding unsupplemented group, ^*p*<0.05, ^^*p*<0.01,^^^*p*<0.001,^^^^*p*<0.0001. ^&^significant between-group difference vs VEH/CON+LR, ^&&&^*p*<0.001, ^&&&&^*p*<0.0001. ^@^significant within group dishabituation of social odor 1 (social 1,3 vs social 2,1), ^@^*p*<0.01-0.0001. Color of statistical symbols represents specific group. Statistical results are summarized in Table 1. *n*, 8-11 subjects/group (A), *n*, 6-12 subjects/group (B)

**Table 1.**
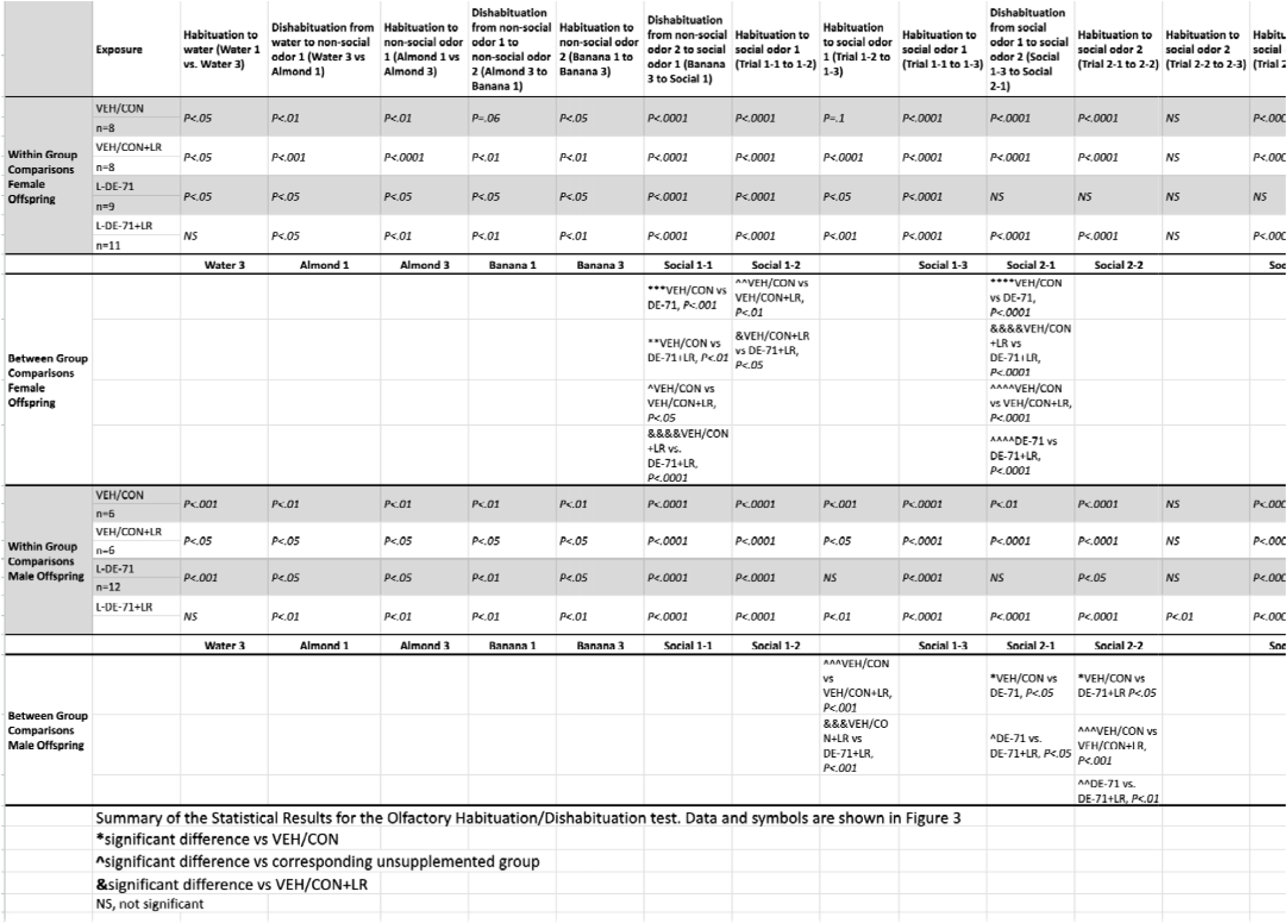
Statistical Results for the Olfactory Habituation/Dishabituation test.

For males, within-group comparisons of habituation for social odors revealed that the DE-71 group behaved differently than the others. DE-71 males were unable to habituate to social odor 1 unlike the other groups (*p*<0.05-0.001) nor did they dishabituate from social odor 1 to 2 in comparison to other groups (*p*<0.01-0.001). Importantly, the DE-71+LR group behaved similarly (habituation) or better (dishabituation social odor 1 to 2, *p*<0.05) than VEH/CON. Also, the LR supplemented VEH/CON group showed improved social odor discrimination as compared to VEH/CON in males (*p*<0.001) and in females (social 1 *p*<0.05, social 2 *p*<0.0001).

### DE-71 decreases pain sensitivity in male offspring; LR Protection

To assess potential alterations in OXT-relevant thermal nociception, we conducted the hot plate test in offspring. Latency to respond was unchanged across treatment groups in females. In contrast, male group comparisons to VEH/CON showed that DE-71 exposed males exhibited significantly increased response latency, indicating reduced thermal sensitivity (*p*<0.01), an effect that was normalized in LR supplemented DE-71 group (*p*<0.05; **Supplementary Fig. 5**).

### DE-71-induced deregulation of *Mct8* and *Dio3* transcript levels in PVH OXTergic neurons; sex-specificity & protection by LR

Having seen protection against socioemotional behavior deficits of DE-71 with LR treatment, that boosts T4 and OXT levels (Sgritta et al., 2019; Varian et al., 2014, 2017), we explored effects of DE-71 exposure and LR treatment on T4 regulatory gene markers in PVH OXT neurons. We used multiplex RNA *in situ* hybridization to quantify *Mct8* and *Dio3* transcripts, in combination with IF of OXTergic neurons. **Figure 4** shows that *Mct8* transcripts on OXT neurons, OXT*^Mct8^*, are upregulated in DE-71 vs VEH/CON females (*p*<0.0001), an effect that is normalized with LR supplementation (vs VEH/CON, ns, **Fig. 4C,E**). Also, LR treatment upregulated OXT*^Mct8^* expression in VEH/CON (*p*<0.05, **Fig. 4D,E**). Similar results were observed in the general (DAPI-positive) cell PVH population (DE-71 vs VEH/CON, *p*<0.0001, **Fig. 4D,F**). *Dio3* transcripts on female OXT neurons, OXT*^Dio3^*, were affected in an opposite direction, i.e., downregulated, in DE-71 (*p*<0.001) and also prevented with LR supplementation (vs DE-71, *p*<0.05, **Fig. 4C,G**). *Dio3* counts were also decreased on all DAPI-positive cells in DE-71 females (*p*<0.0001), and prevented by LR (vs DE-71, *p*<0.01, **Fig. 4D,H**).

**Figure 4.**
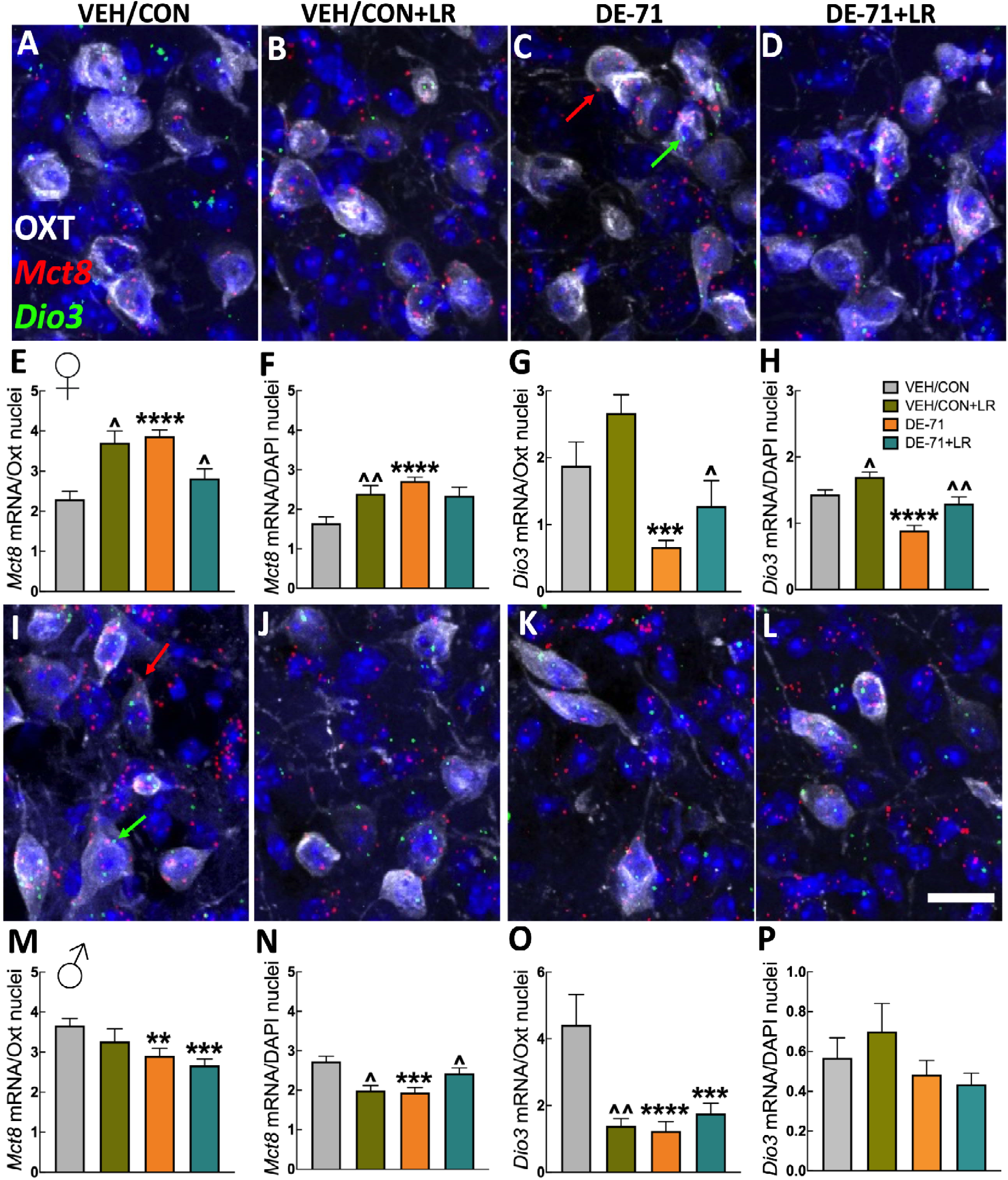
L-DE-71 oppositely deregulates *Mct8* and *Dio3* transcript levels in female and male PVH populations. Representative micrographs of dual IF and multiplex RNA *in situ* hybridization for *Mct8* (red) and *Dio3* mRNA (green) on PVH OXT- and DAPI-positive cells in females (**A-D**) and males (**I-L**)*. Mct8* and *Dio3* mRNA transcripts are homogeneously distributed across the PVH. Mean group values of *Mct8* mRNA transcript counts on OXT neurons (**E**) and on total DAPI-positive cells in females (**F**) and males (**M,N,** respectively). Mean *Dio3* mRNA transcript counts on OXT neurons and total DAPI-positive cells in female (**G,H**) and male (**O,P**, respectively). *statistical difference vs VEH/CON, \*\**p*<0.01, \*\*\**p*<0.001, \*\*\*\**p*<0.0001. **^**statistical difference vs corresponding unsupplemented group, ^*p*<0.05, ^^*p*<0.01. *n*, 3-5 mice/group (E-H); 4-5 mice/group (M-P). Scale bar, 50 μm.

In males, DE-71 reduced Mct8 expression on OXT neurons (*p*<0.01) and DAPI cells (*p*<0.001) as compared to VEH/CON (**Fig. 4K,L,M,N**). LR treatment offered protection from DE-71-induced changes only in DAPI-positive cells expressing *Mct8* (*p*<0.05, **Fig. 4L,N**). PVH OXT*^Dio3^* was decreased in DE-71 (*p*<0.0001) and DE-71+LR (*p*<0.01, **Fig. 4K,L,O**). There were no group differences observed in *Dio3* expression on DAPI cells (**Fig. 4P**).

We also examined effects of DE-71 with and without LR treatment of OXT neuronal count in the PVH. Computer-assisted densitometry indicated significant group differences only in male PVH; DE-71 decreased neuron counts (*p*<0.05, **Supplementary Fig. 6A,B**). LR supplementation had mixed effects, i.e., LR normalized reduction in DE-71-exposed males (ns vs VEH/CON) but reduced OXT neuron counts in controls (*p*<0.05). In summary, the effects of DE-71 include opposite regulation of *Mct8* and *Dio3* on PVH OXT neurons in males and females and LR produces protection in both sexes, albeit to a broader degree in females.

### Female-specific DE-71-induced deregulation of *Mct8* and *Dio3* transcript levels in SON OXTergic neurons; protection by LR

Next, we examined TH regulatory gene markers on SON OXT neurons since these neurons can also regulate social recognition via dendritic projections to the amygdala (Takayanagi et al., 2005). Figure 5 shows raw images of mRNA transcripts after SON sections were processed with dual IF/multiplex RNA in situ hybridization. We found that, in contrast to female PVH, OXT*^Mct8^*expression in female SON is downregulated by DE-71 vs VEH/CON (*p*<0.0001, **Fig. 5C,E**). *Mct8* was also decreased in the general (DAPI-positive) cell population (*p*<0.05, **Fig. 5C,F**). Altered *Mct8* mRNA counts in both SON populations were significantly mitigated by LR: DE-71+LR vs DE-71 for OXT*^Mct8^* (*p*<0.05). Similar results were obtained for female OXT*^Dio3^* (*p<*0.0001, **Fig. 5C,G**). *Dio3* expression in SON DAPI-positive was also downregulated (*p*<0.001) (**Fig. 5C,H**). LR supplementation protected (vs DE-71, *p*<0.05, **Fig. 5D,G**) or normalized (vs VEH/CON, ns, **Fig. 5D,H**) against DE-71 effects in OXT and DAPI cell populations, respectively. In males there were no group effects in *Mct8* and *Dio3* levels in the SON (**Fig. 5I-P**).

**Figure 5.**
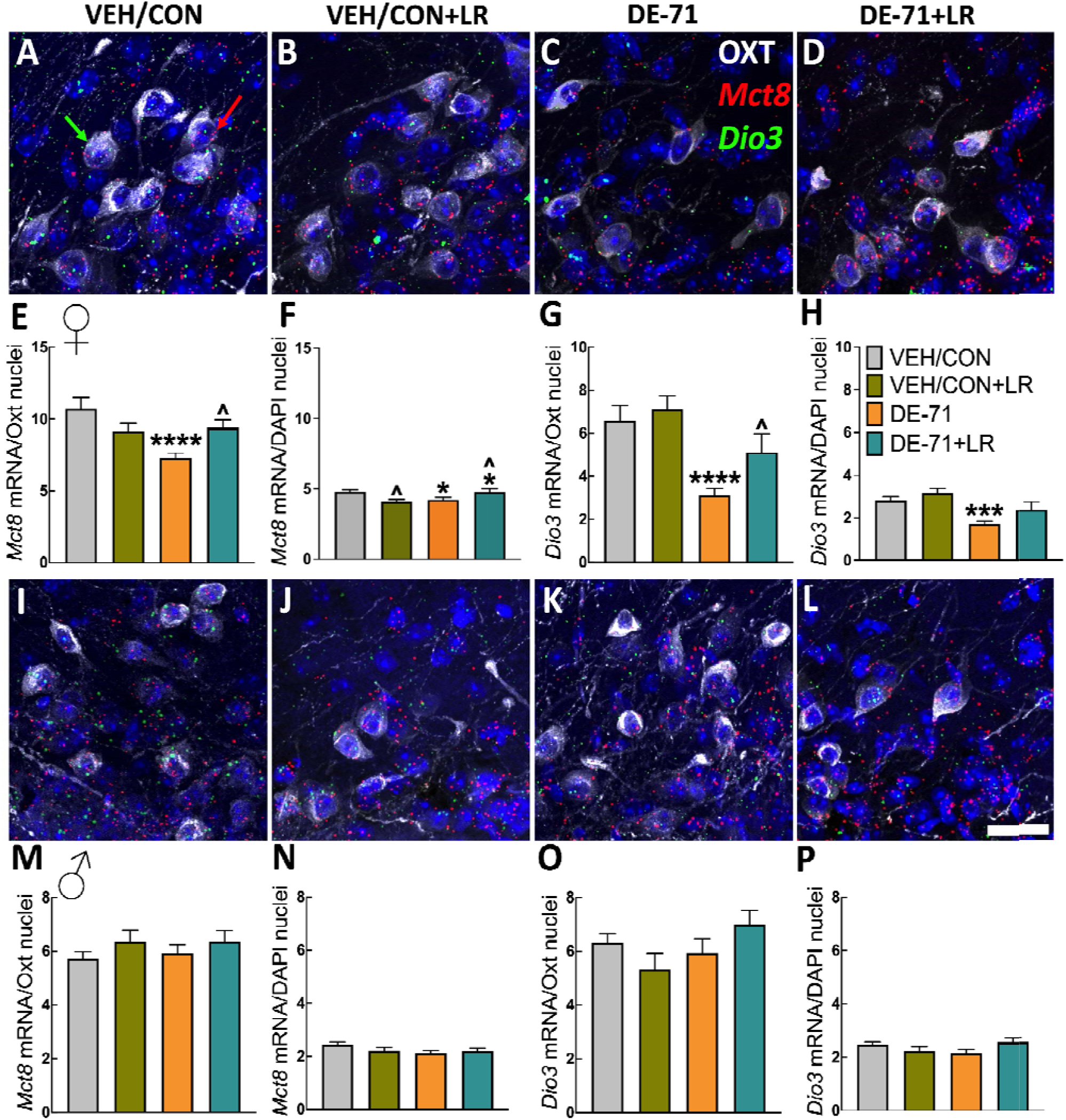
DE-71 deregulation of *Mct8* and *Dio3* transcript levels in SON of females but not males. Representative micrographs of *Mct8* (red) and *Dio3* mRNA (green) expression in OXT immunoreactive neurons and all cells in female SON (**A-D**) and male SON (**I-L**). *Mct8* and *Dio3* mRNA transcripts are homogeneously distributed throughout the SON. Mean group counts of *Mct8* mRNA transcripts on OXT neurons and total DAPI cells in female (**E,F**) and male SON (**M,N**). Mean group counts for *Dio3* mRNA transcripts on OXT neurons and total DAPI-positive cells in females (**G,H**) and males (**O,P**), respectively. *statistical difference vs VEH/CON, \**p*<0.05, \*\*\**p*<0.001, \*\*\*\**p*<0.0001. **^**statistical difference vs corresponding unsupplemented group, ^*p*<0.05. *n*, 3-5 mice/group (E-H); 4-5 mice/group (M-P). Scale bar, 35 μm.

OXT neuronal counts in SON indicated significant changes in DE-71 in females (elevated) and males (reduced) compared to VEH/CON (*p*<0.05). LR supplementation prevented the females (vs DE-71, *p*<0.05) but not male changes (**Supplementary Fig. 6C,D**).

### DE-71 produced no effect on *Oxtr* transcripts in the hippocampus

Since neurons expressing *Oxtr* in the anterior CA2/CA3 subfield of the dorsal hippocampus are implicated in social recognition memory (Raam et al., 2017), we examined the effect of DE-71 on *Oxtr* expression on DAPI+ cells in this region. *Oxtr* was not different in DE-71 groups but decreased in both LR supplemented groups in both sexes (**Supplementary Fig. 7**).

### Sex-dependent dysbiosis in DE-71-exposed offspring; LR protection

Gut dysbiosis is a commonly occurring commorbidity amongst individuals with autism (Fattorusso et al., 2019) and environmental toxicants including PBDE-28 reduce alpha diversity of gut microbiome in infants of mothers with elevated breastmilk burden (Iszatt et al., 2019) and experimental mice (S. Kim et al., 2023). We previously reported less alpha and beta diversity in gut microbiome of mice developmentally exposed to DE-71 with LR treatment providing protection in exposed postnatal offspring (Denys et al., 2025). In this study, we examined the effects of DE-71 and LR in adulthood, at the time of behavioral testing. Using 16S rRNA gene sequencing we first evaluated alpha diversity, a measure of species richness and evenness, represented by Chao1 index scores in the fecal bacterial population. Rarefraction was performed from 0 to 1000 reads to confirm adequate read depth. Mean Chao1 scores shown in Figure 6A demonstrate less microbial richness in female DE-71 vs VEH/CON (*p*<0.0001) and LR supplementation was protective (DE-71+LR vs DE-71, *p*<0.0001). There were no group differences in microbial richness in males (**Fig. 6B**).

**Figure 6.**
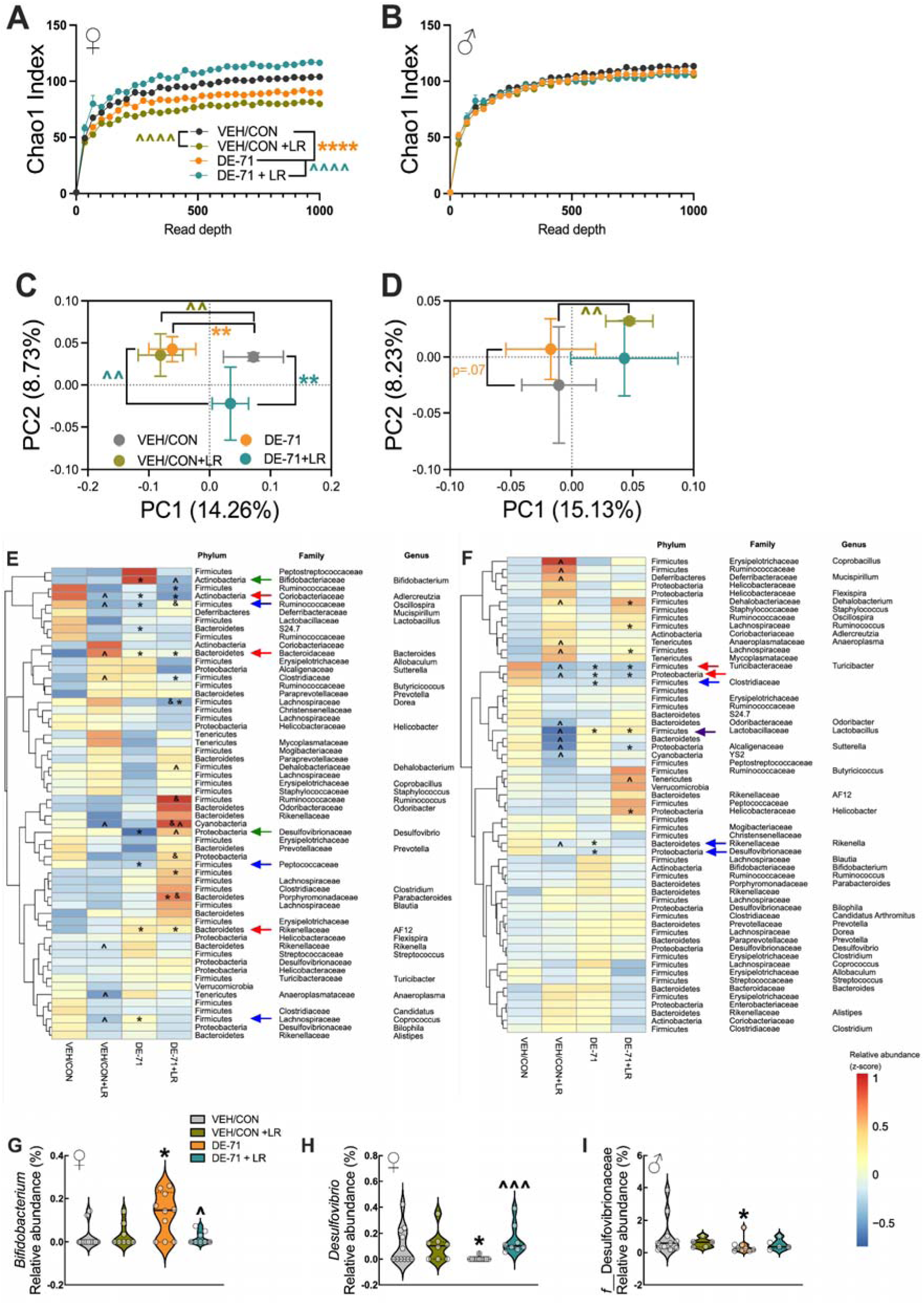
Dysbiosis in adult offspring of dams exposed to DE-71 with or without LR supplementation. **(A)** Adult female Chao1 Index. **(B)** Adult male Chao1 Index. **(C,D)** Principal coordinate analysis (PCoA) plots showing across-treatment variation of the fecal microbiome in female (C) and male offspring (D) using unweighted Unifrac. (**E-F**) Heatmap of most abundant microbial taxa identified by mapping 16S rRNA in female (**E**) and male offspring (**F**). (**G-H**) Relative abundance of selected taxa altered by DE-71 and protected or normalized by additional LR supplementation in females (**I**) Relative abundance of taxa altered by DE-71 in males. *statistical difference vs VEH/CON, \**p*<0.05-0.001; ^statistical difference vs corresponding unsupplemented group, *p*<0.05-0.001. *indicates significant difference vs VEH/CON, **q<0.01; ^ vs corresponding control, ^^q<0.01 using PERMANOVA (**C, D**). **Green** arrows indicate taxa that were altered by DE-71 and normalized by additional LR supplementation. **Blue** arrowheads indicate taxa altered by only in DE-71 but not DE-71+LR vs VEN/CON. **Red** arrows indicate altered by DE-71 and DE-71 + LR. **Purple** arrow indicates group profiles for *Lactobacillus*. Females *n*, 8-11/group; males *n*, 5-10/group

Next, we performed Principal coordinate analysis (PCoA) using unweighted UniFrac distances and PERMANOVA to examine microbial community composition. The most substantial effect on β-diversity was that of DE-71 on female offspring which clustered differently vs VEH/CON (*p*<0.01) along PC1 (14.3%) (**Fig. 6C**). DE-71+LR partially mitigated the effect of DE-71 alone (vs DE-71, *p*<0.01). In males, the DE-71 effect is only *apparent* along PC2 (8.23%) in overall community structure compared to VEH/CON (*p*=0.07). Of note, LR supplemented controls (VEH/CON+LR) clustered similarly with DE-71 (vs VEH/CON, *p*<0.01) in females only, suggesting no benefit offered and potential adverse effect of LR under control conditions. In males LR treatment had an opposite (positive) effect on VEH/CON controls (*p*<0.01, **Fig. 6D**). In summary, results suggest that LR treatment protects against DE-71 effects on alpha-diversity, shown only by females, and in beta-diversity in PBDE-exposed females. In males, exposure and treatment effects on beta-diversity were less salient.

Hierarchical clustering analysis conducted at the genus level identified 75 bacterial taxa in male and female offspring. A heatmap of z-score-transformed relative abundance revealed statistically significant group differences vs VEH/CON in female offspring (**Fig. 6E**). DE-71-exposed females displayed changes in abundance of taxa such as Phylum Actinobacteria Family Bifidobacteriaceae Genus Bifidobacterium (elevated, *p*<0.05, **Fig. 6G**) and Phylum Proteobacteria Family Desulfobrionaceae Genus Desulfovibrio (reduced, *p*<0.05, **Fig. 6H**). These effects were prevented in DE-71+LR (vs. DE-71, *p*<0.05-*p*<0.001). These rescued taxa are indicated with green arrows on the heatmap. Other DE-71 induced alterations included Phylum Firmicutes Family Peptococcaceae (reduced, *p*<0.01), Phylum Firmicutes Family Ruminococcaceae Genus *Oscillospira* (reduced, *p*<0.05) and Phylum Firmicutes Family Lachnospiraceae Genus *Coprococcus* (elevated, *p*<0.05). These taxa were normalized in DE-71+LR (vs VEH/CON, ns) and are indicated with blue arrows on the heatmap. Finally, the following taxa were deregulated by DE-71 and DE-71+LR alike: Phylum Actinobacteria Family Coriobacteriaceae Genus *Aldercreutzia* (reduced, *p*<0.01), Phylum Bacteroidetes Family Bacteroidaceae Genus *Bacteroides* (elevated, *p*<0.01) and Phylum Bacteroidetes Family Rikenellaceae Genus *AF12* (elevated, *p*<0.05). These resistant taxa are indicated with red arrows on the heatmap.

In males, alterations in relative abundance included Phylum Firmicutes Family Clostridiaceae (reduced, *p*<0.05), Phylum Bacteroidetes Family Rikenellaceae Genus *Rikenella* (reduced, *p*<0.05), and Phylum Proteobacteria Family Desulfovibrionaceae (reduced, *p*<0.05, **Fig. 6F**). These alterations were normalized by LR supplementation (vs VEH/CON, ns). These taxa are indicated with blue arrows on the heatmap. When compared to VEH/CON, two taxa in DE-71 and DE-71+LR showed similar reductions in abundance: Phylum Firmicutes Family Turicibacteraceae Genus *Turicibacter* and Phylum Proteobacteria, (*p*<0.05) and similar upregulation: Phylum Firmicutes Family Lactobacillaceae Genus *Lactobacillus* (*p*<0.05).

Of note, DE-71 exposure downregulated the abundance of Family Desulfobrionaceae in both sexes and LR supplementation protected against this effect only in females (**Fig. 6H,I**). LR supplementation upregulated Phylum Firmicutes Family Lactobacillaceae Genus *Lactobacillus* in DE-71 males but downregulated it in VEH/CON males (vs. VEH/CON, *p*<0.05) (**Supplementary Fig. 8**). This taxa was unaffected in female offspring. Yet a different profile for Genus *Lactobacillus* was observed in dams (see below), i.e., DE-71 caused an *apparent* reduction in abundance (vs. VEH/CON, *p*=0.1) while LR treatment mitigated the effects towards normalization (**Fig. 7D**).

**Figure 7.**
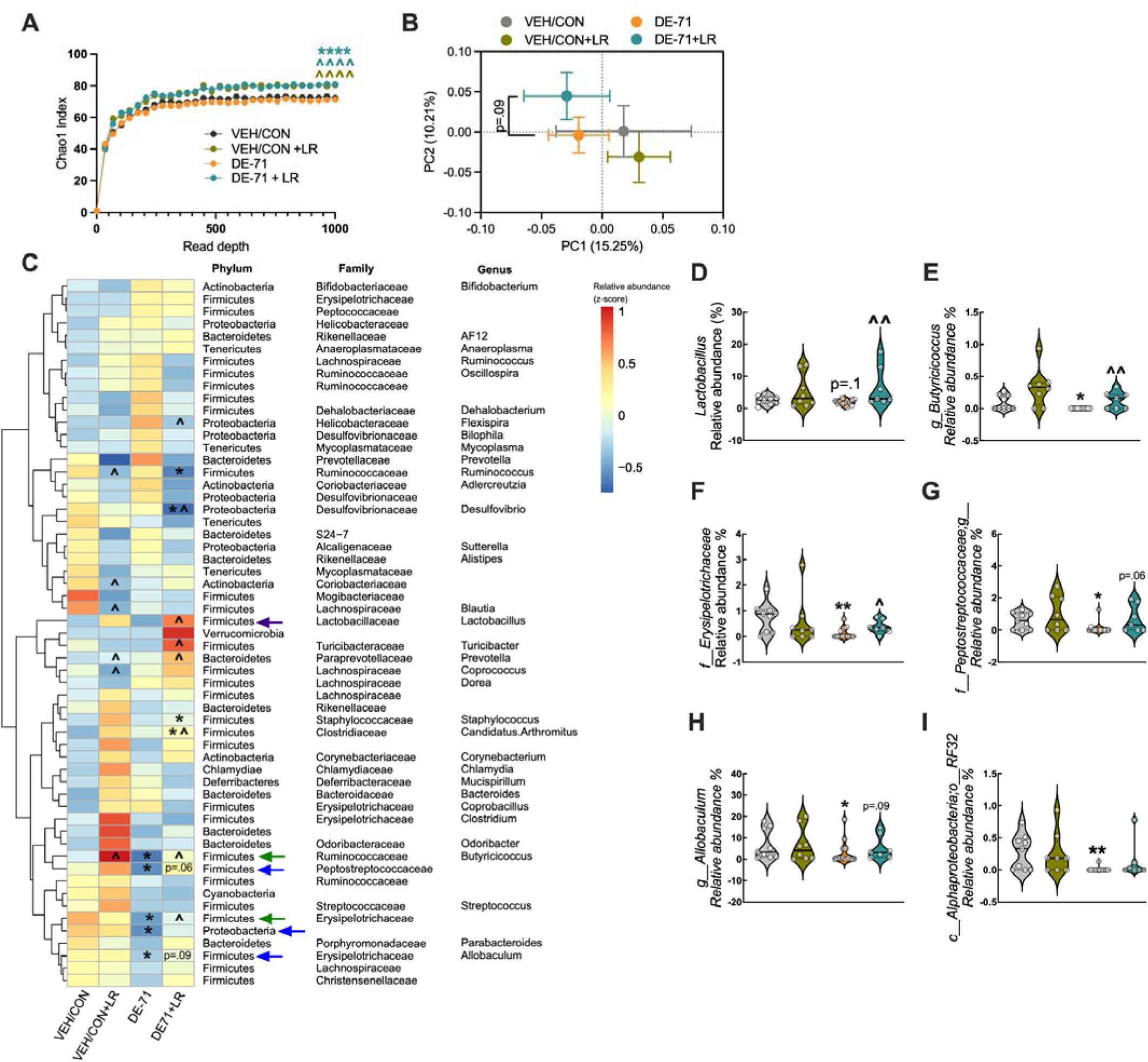
Dysbiosis in dams exposed to DE-71 with or without LR supplementation. **(A)** Chao1 Index. **(B)** Principal coordinate analysis (PCoA) plots showing across-treatment beta-diversity of the fecal microbiome using unweighted UniFrac. (**C**) Heatmap of most abundant microbial taxa. (**D-H**) Relative abundance of taxa altered by DE-71 and/or LR supplementation. *statistical difference vs VEH/CON, \**p*<0.05-0.001; ^statistical difference vs corresponding unsupplemented group, *p*<0.05-0.001. **Green** arrows indicate taxa that were altered by DE-71 and protected by additional LR supplementation. **Blue** arrowheads indicate taxa altered by only in DE-71 but not DE-71+LR vs VEN/CON. **Purple** arrow indicates group profiles for *Lactobacillus*. *n*, 7-12/group.

### Minimal Dysbiosis in dams exposed to DE-71 with or without LR supplementation

Analysis of the 16S rRNA gene sequencing on dam fecal samples indicated no major effects of DE-71 on α-diversity as compared to VEH/CON. However, LR supplemented groups increased α-diversity in VEH/CON (*p*<0.0001) and DE-71 (*p*<0.0001) (**Fig. 7A**). Principal coordinate analysis (PCoA) plots and PERMANOVA revealed no significant effect of DE-71 exposure on β-diversity of gut bacterial community. Only LR supplemented DE-71 samples showed an *apparent* effect along PC2 (10.2%) (q=0.09, **Fig. 7B**).

With regard to genus-level changes, a heatmap of z-score-transformed relative abundance revealed statistically significant group differences vs VEH/CON in female offspring (**Fig. 7C**). DE-71 exposure reduced several taxa in Phyla Firmicutes and Proteobacteria. In particular, DE-71 exposure downregulated Phylum Firmicutes Family Ruminoccoceae Genus *Butyricicoccus* (*p*<0.05, **Fig. 7E**) and Phylum Firmicutes Family Erysipelotrichaceae (*p*<0.01, **Fig. 7F**); effects that were mitigated by LR treatment (vs DE-71, *p*<0.05-0.01). These rescued taxa are indicated with green arrows on the heatmap. Phylum Firmicutes Family Peptostreptococcaceae (*p*<0.05, **Fig. 7G**) Phylum Firmicutes Family Erysipelotrichacceae Genus *Allobaculum* (*p*<0.05, **Fig. 7H**) and Phylum Proteobacteria Class Alphaproteobacteria Order RF32 (*p*<0.01, **Fig. 7I**) were downregulated in DE-71-exposed dams but not in exposed dams supplemented with LR, as compared to VEH/CON, suggesting a beneficial effect of probiotic treatment. These rescued taxa are indicated with blue arrows on the heatmap.

First, LR treatment increased the relative abundance of Phylum Firmicutes Family Lactobacillaceae Genus *Lactobacillus* (DE-71+LR vs DE-71, *p*<0.05, **Fig. 7C**, purple arrow, **Fig. 7D**). In summary, these data indicate minimal effects of adult exposure to DE-71 and some of these changes were protected by LR treatment. As expected, LR supplementation increased the abundance of Genus *Lactobacillus* in the DE-71-exposed dam gut as well as the beneficial changes in alpha and altered beta diversity (apparent).

## Discussion

In accordance with the developmental origins of health and disease (DOHaD) hypothesis, there is increasing evidence that maternal exposure to environmental stressors during the perinatal period may lead to long-term health consequences to offspring during adulthood (D. J. P. Barker, 2007). In previously published reports, we showed that perinatal exposure to the common commercial mixture of Penta-BDEs, DE-71, produced ASD-like social deficits, hypothalamic neuroendocrine disruption and abnormal neurodevelopmental benchmarks in male and female mice offspring (Denys et al., 2025; Kozlova et al., 2022; Kozlova, Gonzalez, et al., 2025). Recently, we also demonstrated that maternal supplementation with LR 6475 probiotic treatment during pregnancy and lactation to DE-71-exposed moms improved developmental benchmarks and normalized adult metabolic dysfunction (Denys et al., 2025). In the current study, we extend our findings by demonstrating that *L. reuteri* treatment protects against PBDE-altered socioemotional recognition and social odor discrimination. We also explore a mechanistic link between developmental DE-71 exposure and reprogrammed TH markers on hypothalamic OXT neurons, and the corrective effects of LR, that may explain DE-71-altered hypothalamic OXT and TH status in ASD-like behavior (Kozlova, Denys, et al., 2025; Kozlova et al., 2022; Kozlova, Gonzalez, et al., 2025), both of which are protected by LR treatment in other mouse models of ASD (Sgritta et al., 2019; Varian et al., 2014, 2017). The potential ability to prevent ASD phenotypes in humans with probiotic therapy is particularly noteworthy because current therapeutic options provide limited benefit and fall short of curing the core symptom domains of ASD.

First, we examined the effects of DE-71 and LR 6475 on maternal health and behavior since these factors can influence offspring outcomes. Neither DE-71 exposure nor maternal LR treatment significantly affected gestational food intake or weight gain or pup retrieval latencies, suggesting that offspring outcomes cannot be attributed to changes in maternal health. Only nest quality was notably improved with maternal LR supplementation in DE-71 dams in agreement with LR-improved maternal care shown by others (Benlakehal et al., 2025; Curley & Champagne, 2016). More importantly, LR supplementation elevated plasma T4 levels in dams exposed to DE-71, consistent with documented T4-promoting actions of LR 6475 (Varian et al., 2014). These results suggest that improved TH functions may serve as a possible mechanism underlying the behavioral, endocrine and molecular disruptions by PBDEs and improvements provided by LR. Similarly, we found that the secondary sex ratio was decreased in LR supplemented groups (yielding greater number of female pups) as reported previously (Erdman, 2014), but this shift is unlikely to account for the beneficial effects of LR treatment on socioemotional behavior and neuromolecular correlates in our offspring. In summary, DE-71 exposure and LR supplementation did not adversely impact basic reproductive parameters, maternal health, maternal behavior.

Next, we examined the effects of perinatal DE-71 exposure and LR treatment on offspring socioemotional behavior. The salient ASD-relevant phenotypes produced by 0.1 mg/kg DE-71 included abnormal SNP and emotional recognition in female and male offspring. Females also showed abnormal SRM and both sexes showed deficient olfactory discrimination. Notably, LR protected against all abnormalities indicating the powerful modulation of social behavioral brain circuits by gut microbiome function and implicating a role for gut-brain signaling. Previously, we reported that LR treatment alleviated exaggerated marble burying in females, but not males (Denys et al., 2025). The sex differences in DE-71 and/or LR effects are likely due to the sexual dimorphism observed in the neurochemistry influencing social recognition ability and repetitive locomotor behavior (Albers, 2012; Gabor et al., 2012; Jiang et al., 2024; Rigney et al., 2024). Our results reflect findings in previous studies that reported restoration of social deficiencies by *L. reuteri* 6475 across ASD mouse models of various etiologies, including monogenetic (*Shank3B*^-/-^), idiopathic (BTBR) and environmental murine models (maternal high-fat diet (HFD), valproic acid, lipopolysaccharide) (Buffington et al., 2016; Mazzone et al., 2024; Sgritta et al., 2019; Wang et al., 2024; Yang et al., 2025).

One component of social cognition is the ability to decipher the emotional expressions of others, a key domain altered in autistic individuals (Frith & Frith, 2007). Recent findings suggest that mice have the ability to discriminate between conspecifics exhibiting altered emotional states (Ferretti et al., 2019). In this study, we discovered that DE-71 significantly reduces emotional recognition ability with LR providing protection in both sexes. This is the first report of exposure to an environmental toxicant altering the emotional cognition domain. Emotional discrimination has been demonstrated to involve OXTergic projections from the PVH to the central amygdala (Ferretti et al., 2019). Thus, it is likely both PBDEs and LR converge on this substrate by oppositely affecting PVH OXT signaling (Kozlova, Denys, et al., 2025). Likewise, the benefits of LR on deficient social behavior have been demonstrated to depend on the OXT receptor (OXTR) (Sgritta et al., 2019). Thus LR may stimulate PVH neurons to release OXT to act at OXTRs in extrahypothalamic brain regions that regulate socioemotional behavior such as the VTA, PFC and amygdala. Although the mechanisms by which *L. reuteri* triggers PVH neurons to increase OXT are not currently known, it may, in part, occur in a vagus nerve-specific manner and the secretion of oxytocin from intestinal epithelia (Danhof et al., 2023; Poutahidis, Kearney, et al., 2013; Sgritta et al., 2019). Together, these findings extend the therapeutic potential of LR by demonstrating its efficacy in preventing socioemotional deficits induced by the environmental toxicant mouse model of ASD.

The parallel effects produced by DE-71 on olfactory signaling may contribute to deficits in socioemotional recognition behavior. Specifically, DE-71 exposed female and male offspring lacked a normal dishabituation response to an initial social odor in comparison to controls, such that they could not distinguish a subsequent new social odor. In addition, DE-71 males showed incomplete habituation to first social odor, in contrast to how control mice function. These results indicate that abnormal social odor discrimination of DE-71-exposed offspring may contribute to social recognition deficits. They may also contribute to emotional recognition abnormalities since emotional recognition ability is dependent on olfactory cues that convey information related to fear emotion of stimulus mice (Ferretti et al., 2019). In our hands, LR prevented both olfactory effects, coincident with protection of DE-71-reduced socioemotional recognition ability, suggesting that both DE-71 and LR act on common substrates. One such possibility is that central OXT and its receptor OXTR within the anterior olfactory nucleus and olfactory cortex and associated regions that contain a high density of *Oxtr* and receives OXT-ergic innervation from the PVN (Knobloch et al., 2012; Oettl et al., 2016). An intriguing hypothesis that can be tested in future studies is that organohalogen toxicants and LR acts on OXTergic social cue processing circuits that have been shown to enhance social recognition/communication (Knobloch et al., 2012; Oettl et al., 2016). Interestingly, LR supplementation of control mice resulted in exaggerated olfactory responses toward social odors relative to VEH/CON in both sexes implicating direct LR activity at olfactory circuits.

In a past clinical trial, the administration of two LR strains (ATCC PTA 6475 and DSM-17938) was tested on children with autism. The treatment improved social functioning but not repetitive behaviors or overall severity of ASD. The benefits of probiotic therapy were ascribed to *L. reuteri* ATCC PTA 6475 since the former but not the latter rescued social deficits but did not affect locomotor activity or increased repetitive behaviors in BTBR mice (Mazzone et al., 2024). Use of *Lactobacillus* treatment has also been used to improve clinical manifestations of neurological impairments not classified under ASD due, in part, to its ability to up-regulate plasma TH and OXT (Poutahidis, Kearney, et al., 2013; Varian et al., 2014, 2017). For example, in certain premature infants with neurological complications, probiotic administration with beneficial bacteria, *Lactobacillus acidophilus* NCDO 1748 and *Bifidobacterium bifidum* NCDO 2203, was associated with improved language scores and reduced severity of neurodevelopmental impairment, but results are mixed and more research is needed to confirm probiotic benefits (Baucells et al., 2023). Like preterm infants, children with autism manifest dysbiosis that includes lower abundance of *Lactobacillus* (one of protective microorganisms along with *Bifidobacterium*) and elevated *Enterobacteriaceae*, including potential pathogenic species such as *Escherichia coli* and *Klebsiella* (Baucells et al., 2023; Darwesh et al., 2024; Fattorusso et al., 2019). These gut microbial alterations have also been seen in some, but not all, mouse ASD models, suggesting that additional or alternative processes related to gut dysbiosis may underlie the neurobehavioral benefits of *Limosilactobacillus reuteri*.

Socioemotional behavior is critically dependent on the central OXT system and a unifying feature of ASD mouse models of various etiology is dysregulated hypothalamic OXT (Buffington et al., 2016; Kozlova, Denys, et al., 2025; Sgritta et al., 2019; Wagner & Harony-Nicolas, 2018). Therefore, we examined whether DE-71 exposure reprograms thyroid hormone (TH)-related neuromolecular regulators in hypothalamic OXT neurons and whether LR restores these changes, given its ability to normalize PVH OXT function in several ASD models. PVH neurons are the likely source of OXT release to activate OXTRs in extrahypothalamic brain regions that regulate socioemotional behavior such as the VTA, PFC and amygdala (Ferretti et al., 2019; Tan et al., 2019). We found that DE-71 exposure deregulated OXT neurons in PVH and SON by reprogramming their expression of *Mct8* and *Dio3* mRNA transcripts, thereby potentially influencing local TH status and brain OXT transcription and signaling (Adan et al., 1992; Nomura et al., 2002; Vasudevan et al., 2001). Sex-dependent alterations included upregulated *Mct8* with downregulated *Dio3* transcripts in female offspring. Reduced Dio3 expression, which would be expected to decrease TH inactivation, together with increased Mct8 expression, which facilitates cellular TH uptake, is consistent with a compensatory increase in TH availability and signaling in OXTergic neurons following PBDE-induced disruption of TH homeostasis. In contrast, male PVH OXT neurons and SON OXT neurons of females expressed downregulated *Mct8* and *Dio3* transcripts, thereby exhibiting reduced TH degradation but having reduced access to THs. Importantly, LR supplementation provided protection against all DE-71-induced deregulation, primarily in females, consistent with a TH-promoting effect of LR on paraventricular hypothalamic OXT circuits involved in social recognition. These results suggest that a) PVH and SON OXT neurons are targets of DE-71 exposure and LR treatment and b) that distinct sex-specific TH regulatory neuromolecular mechanisms on hypothalamic OXT neurons may underlie the abnormal socioemotional behavior in DE-71 exposed offspring. These novel findings complement reports by Varian and team (2017) of increased plasma OXT after administration of LR 6475 lysates to adult female mice (Varian et al., 2017) and normalized brain OXT in male offspring of LR supplemented HFD dams (Buffington et al., 2016). The evidence we present here, of the LR-rescuable, deficient TH status of PVH OXT neurons, in combination with ASD-like phenotypic behavior produced by PBDEs, supports the hypothesis that LR mitigates toxicant-induced ASD-like traits by converging on oxytocin and thyroid hormone signaling pathways.

As gut microbial dysbiosis is a common feature of ASD and may contribute to abnormal social behavior, we investigated whether perinatal DE-71 exposure altered the gut microbiome and whether these changes were reversed by LR supplementation. 16S rRNA gene sequencing of fecal samples revealed sex-specific alterations in bacterial community structure in adult offspring. An examination of gut bacterial community structure using 16S rRNA gene sequencing on fecal samples revealed sex-specific alterations caused by DE-71 exposure in adult offspring. LR normalized the DE-71-induced reduction in - and -diversity only in females, effects that were prevented by LR supplementation. Alterations in taxa-level abundance caused by DE-71, and removed by LR, were observed in both sexes. These involved bacteria such as beneficial ones, i.e., *Bifidobacterium, Coprococcus, Oscillospira*, and conditionally harmful, i.e., Peptococcaceae in females. Beneficial bacteria, *Rikenella*, and *Turicibacter,* were also altered in males. Moreover, downregulation of taxa in Phylum Proteobacteria Family Desulfovibrionaceae that was observed in both DE-71-exposed females and males may be disadvantageous since bioremediation using microbial tools such as *Desulfovibrio* spp. (and *Dehalococcoides*) have the potential for deactivating PBDEs in the environment via anaerobic debromination but whether this function contributes to PBDE metabolism in the mammalian gut remains unknown. (Y.-M. Kim et al., 2007; Robrock et al., 2009). LR supplementation protected against this reduction in exposed female, but not male offspring. We speculate that the decreased relative abundance of *Desulfovibrio* may augment the effects of PBDEs *in vivo*, and that LR may provide benefit in a sex-specific manner. The significant rise in Phylum Actinobacteria Family Bifidobacteriaceae Genus *Bifidobacterium,* a beneficial bacterium, in DE-71-exposed female offspring likely represents a compensatory process to toxicant exposure whereby *Bifidobacterium* can rapidly multiply to occupy the resources and space left behind by more sensitive beneficial bacteria like *Lactobacillus*. *Bifidobacterium* in the gut provides defense against toxic insults by binding, sequestering, or actively metabolizing environmental chemicals (Teffera et al., 2024). Notably, LR treated offspring and dams showed augmented relative abundance in *Lactobacillus* in fecal samples through P40 but not with continued treatment. Nevertheless, others have shown that the benefits of LR can occur both in the presence and absence of *L. reuteri* colonization or establishment in the gut (Haileselassie et al., 2016; Poutahidis, Kleinewietfeld, et al., 2013). These LR benefits can occur by strengthening the gut barrier, regulating the immune system, and shifting the balance towards beneficial bacteria in the gut through its production of metabolites like reuterin, organic acids, and ethanol (Ciorba, 2012; Peng et al., 2023). Work by Poutahidis and team (2013) has demonstrated that LR can promote OXT secretion in a vagus nerve-specific manner, suggesting a mechanism involving gut-brain signaling (Mu et al., 2018; Poutahidis, Kearney, et al., 2013).

## Conclusions

Our findings show that developmental exposure to PBDE toxicants may facilitate an autistic phenotype by reprogramming socioemotional behavior and olfactory social odor discrimination ability. Another novelty of our findings pertains to the altered neuromolecular TH-regulatory markers on hypothalamic OXT neurons suggesting oxytocin-thyroid signaling crosstalk is involved in this reprogramming. Toxicity of OXT-thyroid signaling warrants more study since disrupted social behavior occurs, in conjunction with an altered hypothalamic OXTergic profile as a result of exposure to other thyroid-disrupting chemicals such as Bisphenol A (BPA), chlorpyrifos, Dichlorodiphenyltrichloroethane (DDT), genistein, and polychlorinated biphenyls (PCBs) (Gore et al., 2019, 2023; Patisaul, 2017). Finally, we show protection from abnormalities in behavior, olfactory processing and neuromolecular regulation by LR supplementation, indicating an important role of microbiome-gut-brain signaling in neurodevelopmental alterations caused by early life toxicant burden. These findings support the TH- and OXT-promoting reports of LR treatment reported by others. This extends from our previous report that DE-71-induced reduction in maternal levels of T4 and altered offspring developmental benchmarks are mitigated by LR (Denys et al., 2025; Kozlova, Denys, et al., 2025). In addition, LR may sway the balance within the gut microbiome to xenobiotic biotransforming bacteria (Clarke et al., 2019; Ju et al., 2019) or act as a direct biodetoxification agent in addition to providing other benefits. In conclusion, our study shows that probiotic therapy may provide therapeutic protection against ASD-relevant neurodevelopmental and behavioral reprogramming by environmental pollutant exposure during early life. Thus, leveraging the gut microbiome to combat autistic-like deficits may be worthy of further investigation.

## Acknowledgments

We thank Dr. Harold Gainer (deceased) for gift of PS38 Neurophysin antibody. We are grateful to Drs. zur Nieden and Sladek (UC Riverside) lab for the gift of mice. We thank L. Campoy and Y. Korde for technical assistance. We thank Drs. T. Garland (UCR) and Hartman (Loma Linda University) for the use of Echo MRI and Ethovision software, respectively. We thank Drs. D. Carter (UCR), J. Lopez and A. Gilman (Evident Scientific) for assistance with imaging. We are grateful to Drs. Levi Maston and Scott Rowlins (Indica Labs) for help with HALO AI software. Images were created with Biorender.com.

## Declarations

### Disclaimer

Research reported in this publication was supported by the National Institute Of Environmental Health Sciences of the National Institutes of Health under Award Numbers F31ES034304 and F31AI179030. The content is solely the responsibility of the authors and does not necessarily represent the official views of the National Institutes of Health.

### Ethics approval

Care and treatment of animals was performed in accordance with guidelines from and approved by the University of California, Riverside Institutional Animal Care and Use Committee (AUP# 5 and 20210031).

### Consent to participate

Not applicable.

### Consent for publication

All authors reviewed and approved the final manuscript.

### Permission to reproduce material from other sources

Two of the seven data bars in one panel of figure 1 (Fig. 1I) contains male plasma oxytocin data that also appears in a related manuscript currently under review. The data are reused with permission of the authors, as they originate from the same research program and have not yet been published elsewhere. The figure that shares these 17 data points contains additional measures from additional animals to produce a better estimate of the effect size of L-DE-71 exposure (only in males). In our opinion, this limited amount of data overlap does not constitute dual publication.

### Data availability

The 16S rRNA gene sequences have been deposited in the National Center for Biotechnology Information (NCBI)’s Sequence Read Archive (SRA) under the SRA BioProject

Accession PRJNA1162038

### Author Contributions

**Conceptualization,** E.V.K., M.E.D., M.C.-C.; **Methodology**, E.V.K., M.E.D., E.A.D., R.L., V.P., A.H., M.C.-C.; **Validation**, E.V.K., M.E.D., A.E.B., E.A.D., R.L., C.N.L., A.L., V.P., A.H., M.C.-C.; **Formal Analysis**, E.V.K., M.E.D., A.E.B., E.A.D., R.L., C.N.L., A.L., V.P., A.H., M.C.-C.; **Investigation**, E.V.K., M.E.D., A.E.B., E.A.D., R.L., C.N.L., A.L., V.P., A.H., M.C.-C.; **Writing – Original Draft**, E.V.K., M.E.D., M.C.-C.; **Writing – Reviewing and Editing**, E.V.K., M.E.D., A.E.B., E.A.D., R.L., C.N.L., A.L., V.P., A.H., M.C.-C.; **Visualization**, E.V.K., R.L.; **Resources**, E.V.K., M.E.D., A.H., M.C.-C.; **Data Curation**, E.V.K., M.E.D., A.E.B., E.A.D., R.L., V.P., A.H., M.C.-C.; **Supervision**, E.V.K., A.H., M.C.-C.; **Project Administration**, E.V.K., M.E.D., A.H., M.C.-C.; **Funding Acquisition**, E.V.K., M.E.D., A.H., M.C.-C.

## Abbreviations

AON: anterior olfactory nucleus
ASD: autism spectrum disorder
AVP: arginine vasopressin
BTBR: Black and Tan Brachyury (inbred mouse strain)
CFU: colony forming units
CNS: central nervous system
CON: control
DAPI: 4’,6-diamidino-2-phenylindole
DE-71: commercial penta-brominated diphenyl ether mixture
DEPC: diethyl pyrocarbonate
*Dio3*: deiodinase iodothyronine type III gene
DOHaD: developmental origins of health and disease
ERT: emotional recognition test
ELISA: enzyme-linked immunosorbent assay
GI: gastrointestinal
HFD: high-fat diet
IF: immunofluorescence
IGF-1: insulin-like growth factor 1
LPS: lipopolysaccharide
LR: *Limosilactobacillus reuteri*
LOS: late-onset sepsis
*Mct8*: monocarboxylate transporter 8 (Slc16a2) gene
mPFC: medial prefrontal cortex
MRS: De Man, Rogosa, and Sharpe (medium)
NA: numerical aperture
NDD: neurodevelopmental disorder
NEC: necrotizing enterocolitis
NMR: nuclear magnetic resonance
OD: optical density
OHT: olfactory habituation/dishabituation test
OXT: oxytocin
*Oxtr*: oxytocin receptor gene
OTU: operational taxonomic unit
PBS: phosphate-buffered saline
PB: phosphate buffer
PBDE: polybrominated diphenyl ether
PCoA: principal component analysis
PCR: polymerase chain reaction
P: postnatal day
PFC: prefrontal cortex
PVH: paraventricular hypothalamic nucleus
QMR: quantitative magnetic resonance
qPCR: quantitative polymerase chain reaction
rRNA: ribosomal ribonucleic acid
RM: repeated measures
ROI: region of interest
RT: room temperature
rT3: reverse triiodothyronine
SEM: standard error of the mean
SNP: social novelty preference
SON: supraoptic nucleus
SRM: social recognition memory
SRMT: social recognition memory test
T3: triiodothyronine
T4: thyroxine
TH: thyroid hormone
µL: microliter
VEH/CON: vehicle control
VPA: valproic acid
VTA: ventral tegmental area

## References

Abbasi, G., Li, L., & Breivik, K. (2019). Global Historical Stocks and Emissions of PBDEs. Environmental Science & Technology, 53(11), 6330–6340. 10.1021/acs.est.8b07032

Adan, R. A., Cox, J. J., van Kats, J. P., & Burbach, J. P. (1992). Thyroid hormone regulates the oxytocin gene. The Journal of Biological Chemistry, 267(6), 3771–3777. 10.1016/S0021-9258(19)50592-0

Alavi, S., Mitchell, J. D., Cho, J. Y., Liu, R., Macbeth, J. C., & Hsiao, A. (2020). Interpersonal Gut Microbiome Variation Drives Susceptibility and Resistance to Cholera Infection. Cell, 181(7), 1533–1546.e13. 10.1016/j.cell.2020.05.036

Albers, H. E. (2012). The regulation of social recognition, social communication and aggression: vasopressin in the social behavior neural network. Hormones and Behavior, 61(3), 283–292. 10.1016/j.yhbeh.2011.10.007

American Psychiatric Association. (2013). *Diagnostic and Statistical Manual of Mental Disorders (DSM-5®)*. American Psychiatric Publishing. https://play.google.com/store/books/details?id=-JivBAAAQBAJ

Audunsdottir, K., Sartorius, A. M., Kang, H., Glaser, B. D., Boen, R., Nærland, T., Alaerts, K., Kildal, E. S. M., Westlye, L. T., Andreassen, O. A., & Quintana, D. S. (2024). The effects of oxytocin administration on social and routinized behaviors in autism: A preregistered systematic review and meta-analysis. Psychoneuroendocrinology, 167(107067), 107067. 10.1016/j.psyneuen.2024.107067

Bankhead, P., Loughrey, M. B., Fernández, J. A., Dombrowski, Y., McArt, D. G., Dunne, P. D., McQuaid, S., Gray, R. T., Murray, L. J., Coleman, H. G., James, J. A., Salto-Tellez, M., & Hamilton, P. W. (2017). QuPath: Open source software for digital pathology image analysis. Scientific Reports, 7(1), 16878. 10.1038/s41598-017-17204-5

Baribeau, D., & Anagnostou, E. (2022). Novel treatments for autism spectrum disorder based on genomics and systems biology. Pharmacology & Therapeutics, 230(107939), 107939. 10.1016/j.pharmthera.2021.107939

Barker, D. J. (1995). Fetal origins of coronary heart disease. BMJ, 311(6998), 171–174. 10.1136/bmj.311.6998.171

Barker, D. J. P. (2007). The origins of the developmental origins theory. Journal of Internal Medicine, 261(5), 412–417. 10.1111/j.1365-2796.2007.01809.x

Baucells, B. J., Sebastiani, G., Herrero-Aizpurua, L., Andreu-Fernández, V., Navarro-Tapia, E., García-Algar, O., & Figueras-Aloy, J. (2023). Effectiveness of a probiotic combination on the neurodevelopment of the very premature infant. Scientific Reports, 13(1), 10344. 10.1038/s41598-023-37393-6

Ben-Barak, Y., Russell, J. T., Whitnall, M. H., Ozato, K., & Gainer, H. (1985). Neurophysin in the hypothalamo-neurohypophysial system. I. Production and characterization of monoclonal antibodies. The Journal of Neuroscience: The Official Journal of the Society for Neuroscience, 5(1), 81–97. 10.1523/JNEUROSCI.05-01-00081.1985

Benlakehal, R., Gaetano, A., Charlet, R., Bouwalerh, H., Cogez, V., Harduin-Lepers, A., Sendid, B., Nicoletti, F., Maccari, S., & Morley-Fletcher, S. (2025). Probiotic L. reuteri treatment in stressed rats improves maternal care and corrects corticosterone-BDNF-oxytocin balance offering insights for maternal postpartum distress treatment. Scientific Reports, 15(1), 23544. 10.1038/s41598-025-05848-7

Bielsky, I. F., Hu, S.-B., Szegda, K. L., Westphal, H., & Young, L. J. (2004). Profound impairment in social recognition and reduction in anxiety-like behavior in vasopressin V1a receptor knockout mice. Neuropsychopharmacology: Official Publication of the American College of Neuropsychopharmacology, 29(3), 483–493. 10.1038/sj.npp.1300360

Bignami, G. (1996). Economical test methods for developmental neurobehavioral toxicity. Environmental Health Perspectives, 104 *Suppl 2*, 285–298. 10.1289/ehp.96104s2285

Braun, J. M., Kalkbrenner, A. E., Just, A. C., Yolton, K., Calafat, A. M., Sjödin, A., Hauser, R., Webster, G. M., Chen, A., & Lanphear, B. P. (2014). Gestational exposure to endocrine-disrupting chemicals and reciprocal social, repetitive, and stereotypic behaviors in 4- and 5-year-old children: the HOME study. Environmental Health Perspectives, 122(5), 513–520. 10.1289/ehp.1307261

Buffington, S. A., Di Prisco, G. V., Auchtung, T. A., Ajami, N. J., Petrosino, J. F., & Costa-Mattioli, M. (2016). Microbial Reconstitution Reverses Maternal Diet-Induced Social and Synaptic Deficits in Offspring. Cell, 165(7), 1762–1775. 10.1016/j.cell.2016.06.001

Caporaso, J. G., Kuczynski, J., Stombaugh, J., Bittinger, K., Bushman, F. D., Costello, E. K., Fierer, N., Peña, A. G., Goodrich, J. K., Gordon, J. I., Huttley, G. A., Kelley, S. T., Knights, D., Koenig, J. E., Ley, R. E., Lozupone, C. A., McDonald, D., Muegge, B. D., Pirrung, M., … Knight, R. (2010). QIIME allows analysis of high-throughput community sequencing data. Nature Methods, 7(5), 335–336. 10.1038/nmeth.f.303

Ciorba, M. A. (2012). A gastroenterologist’s guide to probiotics. Clinical Gastroenterology and Hepatology: The Official Clinical Practice Journal of the American Gastroenterological Association, 10(9), 960–968. 10.1016/j.cgh.2012.03.024

Clarke, G., Sandhu, K. V., Griffin, B. T., Dinan, T. G., Cryan, J. F., & Hyland, N. P. (2019). Gut Reactions: Breaking Down Xenobiotic-Microbiome Interactions. Pharmacological Reviews, 71(2), 198–224. 10.1124/pr.118.015768

Costa, L. G., & Giordano, G. (2007). Developmental neurotoxicity of polybrominated diphenyl ether (PBDE) flame retardants. Neurotoxicology, 28(6), 1047–1067. 10.1016/j.neuro.2007.08.007

Cuesta-Marti, C., Uhlig, F., Muguerza, B., Hyland, N., Clarke, G., & Schellekens, H. (2023). Microbes, oxytocin and stress: Converging players regulating eating behavior. Journal of Neuroendocrinology, 35(9), e13243. 10.1111/jne.13243

Curley, J. P., & Champagne, F. A. (2016). Influence of maternal care on the developing brain: Mechanisms, temporal dynamics and sensitive periods. Frontiers in Neuroendocrinology, 40, 52–66. 10.1016/j.yfrne.2015.11.001

Danhof, H. A., Lee, J., Thapa, A., Britton, R. A., & Di Rienzi, S. C. (2023). Microbial stimulation of oxytocin release from the intestinal epithelium via secretin signaling. In bioRxivorg. 10.1101/2023.03.09.531917

Darwesh, M.-A. K., Bakr, W., Omar, T. E. I., El-Kholy, M. A., & Azzam, N. F. (2024). Unraveling the relative abundance of psychobiotic bacteria in children with Autism Spectrum Disorder. Scientific Reports, 14(1), 24321. 10.1038/s41598-024-72962-3

Denys, M. E., Kozlova, E. V., Liu, R., Bishay, A. E., Do, E. A., Piamthai, V., Korde, Y. V., Luna, C. N., Lam, A. A., Hsiao, A., & Currás-Collazo, M. (2025). Maternal probiotic supplementation protects against PBDE-induced developmental, behavior and metabolic reprogramming in a sexually dimorphic manner: Role of gut microbiome. Archives of Toxicology, 99(1), 423–446. 10.1007/s00204-024-03882-4

Dinan, T. G., Stanton, C., & Cryan, J. F. (2013). Psychobiotics: a novel class of psychotropic. Biological Psychiatry, 74(10), 720–726. 10.1016/j.biopsych.2013.05.001

Ding, G., Yu, J., Cui, C., Chen, L., Gao, Y., Wang, C., Zhou, Y., & Tian, Y. (2015). Association between prenatal exposure to polybrominated diphenyl ethers and young children’s neurodevelopment in China. Environmental Research, 142, 104–111. 10.1016/j.envres.2015.06.008

Drage, D. S., Heffernan, A. L., Cunningham, T. K., Aylward, L. L., Mueller, J. F., Sathyapalan, T., & Atkin, S. L. (2019). Serum measures of hexabromocyclododecane (HBCDD) and polybrominated diphenyl ethers (PBDEs) in reproductive-aged women in the United Kingdom. Environmental Research, 177, 108631. 10.1016/j.envres.2019.108631

Erdman, S. (2014). Maternal gut microbes control offspring sex and survival. Journal of Probiotics & Health, 02(01), 1–6. 10.4172/2329-8901.1000120

Fattorusso, A., Di Genova, L., Dell’Isola, G. B., Mencaroni, E., & Esposito, S. (2019). Autism spectrum disorders and the gut Microbiota. Nutrients, 11(3), 521. 10.3390/nu11030521

Faul, F., Erdfelder, E., Lang, A.-G., & Buchner, A. (2007). G*Power 3: a flexible statistical power analysis program for the social, behavioral, and biomedical sciences. Behavior Research Methods, 39(2), 175–191. 10.3758/bf03193146

Ferretti, V., Maltese, F., Contarini, G., Nigro, M., Bonavia, A., Huang, H., Gigliucci, V., Morelli, G., Scheggia, D., Managò, F., Castellani, G., Lefevre, A., Cancedda, L., Chini, B., Grinevich, V., & Papaleo, F. (2019). Oxytocin signaling in the central amygdala modulates emotion discrimination in mice. Current Biology: CB, 29(12), 1938–1953.e6. 10.1016/j.cub.2019.04.070

Friard, O., & Gamba, M. (2016). BORIS: a free, versatile open-source event-logging software for video/audio coding and live observations. Methods in Ecology and Evolution / British Ecological Society, 7(11), 1325–1330. 10.1111/2041-210x.12584

Friedrich, N., Thuesen, B., Jørgensen, T., Juul, A., Spielhagen, C., Wallaschofksi, H., & Linneberg, A. (2012). The association between IGF-I and insulin resistance: a general population study in Danish adults. Diabetes Care, 35(4), 768–773. 10.2337/dc11-1833

Frith, C. D., & Frith, U. (2007). Social cognition in humans. Current Biology: CB, 17(16), R724–R732. 10.1016/j.cub.2007.05.068

Gabor, C. S., Phan, A., Clipperton-Allen, A. E., Kavaliers, M., & Choleris, E. (2012). Interplay of oxytocin, vasopressin, and sex hormones in the regulation of social recognition. Behavioral Neuroscience, 126(1), 97–109. 10.1037/a0026464

Gascon, M., Vrijheid, M., Martínez, D., Forns, J., Grimalt, J. O., Torrent, M., & Sunyer, J. (2011). Effects of pre and postnatal exposure to low levels of polybromodiphenyl ethers on neurodevelopment and thyroid hormone levels at 4 years of age. Environment International, 37(3), 605–611. 10.1016/j.envint.2010.12.005

Gibson, E. A., Siegel, E. L., Eniola, F., Herbstman, J. B., & Factor-Litvak, P. (2018). Effects of Polybrominated Diphenyl Ethers on Child Cognitive, Behavioral, and Motor Development. International Journal of Environmental Research and Public Health, 15(8). 10.3390/ijerph15081636

Gillera, S. E. A., Marinello, W. P., Horman, B. M., Phillips, A. L., Ruis, M. T., Stapleton, H. M., Reif, D. M., & Patisaul, H. B. (2020). Sex-specific effects of perinatal FireMaster® 550 (FM 550) exposure on socioemotional behavior in prairie voles. Neurotoxicology and Teratology, 79(106840), 106840. 10.1016/j.ntt.2019.106840

Gore, A. C., Krishnan, K., & Reilly, M. P. (2019). Endocrine-disrupting chemicals: Effects on neuroendocrine systems and the neurobiology of social behavior. Hormones and Behavior, 111, 7–22. 10.1016/j.yhbeh.2018.11.006

Gore, A. C., Zoeller, R. T., & Currás-Collazo, M. (2023). Neuroendocrine effects of polychlorinated biphenyls (PCBs). In Advances in Neurotoxicology (pp. 81–135). Elsevier. 10.1016/bs.ant.2023.08.003

Haileselassie, Y., Navis, M., Vu, N., Qazi, K. R., Rethi, B., & Sverremark-Ekström, E. (2016). Postbiotic modulation of retinoic acid imprinted mucosal-like dendritic cells by probiotic Lactobacillus reuteri 17938 in vitro. Frontiers in Immunology, 7, 96. 10.3389/fimmu.2016.00096

Han, X., Meng, L., Li, Y., Li, A., Turyk, M. E., Yang, R., Wang, P., Xiao, K., Zhao, J., Zhang, J., Zhang, Q., & Jiang, G. (2020). Associations between the exposure to persistent organic pollutants and type 2 diabetes in East China: A case-control study. Chemosphere, 241, 125030. 10.1016/j.chemosphere.2019.125030

Heindel, J. J., & Vandenberg, L. N. (2015). Developmental origins of health and disease: a paradigm for understanding disease cause and prevention. Current Opinion in Pediatrics, 27(2), 248–253. 10.1097/MOP.0000000000000191

Herbstman, J. B., Sjödin, A., Kurzon, M., Lederman, S. A., Jones, R. S., Rauh, V., Needham, L. L., Tang, D., Niedzwiecki, M., Wang, R. Y., & Perera, F. (2010). Prenatal exposure to PBDEs and neurodevelopment. Environmental Health Perspectives, 118(5), 712–719. 10.1289/ehp.0901340

Hertz-Picciotto, I., & Delwiche, L. (2009). The rise in autism and the role of age at diagnosis. Epidemiology, 20(1), 84–90. 10.1097/EDE.0b013e3181902d15

Hess, S. E., Rohr, S., Dufour, B. D., Gaskill, B. N., Pajor, E. A., & Garner, J. P. (2008). Home improvement: C57BL/6J mice given more naturalistic nesting materials build better nests. Journal of the American Association for Laboratory Animal Science: JAALAS, 47(6), 25–31. https://www.ncbi.nlm.nih.gov/pubmed/19049249

Hirota, T., & King, B. H. (2023). Autism spectrum disorder: A review. The Journal of the American Medical Association, 329(2), 157–168. 10.1001/jama.2022.23661

Hsiao, E. Y., McBride, S. W., Hsien, S., Sharon, G., Hyde, E. R., McCue, T., Codelli, J. A., Chow, J., Reisman, S. E., Petrosino, J. F., Patterson, P. H., & Mazmanian, S. K. (2013). Microbiota modulate behavioral and physiological abnormalities associated with neurodevelopmental disorders. Cell, 155(7), 1451–1463. 10.1016/j.cell.2013.11.024

Huang, Y., Huang, X., Ebstein, R. P., & Yu, R. (2021). Intranasal oxytocin in the treatment of autism spectrum disorders: A multilevel meta-analysis. Neuroscience and Biobehavioral Reviews, 122, 18–27. 10.1016/j.neubiorev.2020.12.028

Iszatt, N., Janssen, S., Lenters, V., Dahl, C., Stigum, H., Knight, R., Mandal, S., Peddada, S., González, A., Midtvedt, T., & Eggesbø, M. (2019). Environmental toxicants in breast milk of Norwegian mothers and gut bacteria composition and metabolites in their infants at 1 month. Microbiome, 7(1), 34. 10.1186/s40168-019-0645-2

Jiang, X., Sun, M., Chen, J., & Pan, Y. (2024). Sex-specific and state-dependent neuromodulation regulates male and female locomotion and sexual behaviors. Research (Washington, D.C.), 7, 0321. 10.34133/research.0321

Ju, J., Shen, L., Xie, Y., Yu, H., Guo, Y., Cheng, Y., Qian, H., & Yao, W. (2019). Degradation potential of bisphenol A by Lactobacillus reuteri. LWT, 106, 7–14. 10.1016/j.lwt.2019.02.022

Khachadourian, V., Mahjani, B., Sandin, S., Kolevzon, A., Buxbaum, J. D., Reichenberg, A., & Janecka, M. (2023). Comorbidities in autism spectrum disorder and their etiologies. Translational Psychiatry, 13(1), 71. 10.1038/s41398-023-02374-w

Kim, B., Colon, E., Chawla, S., Vandenberg, L. N., & Suvorov, A. (2015). Endocrine disruptors alter social behaviors and indirectly influence social hierarchies via changes in body weight. Environmental Health: A Global Access Science Source, 14, 64. 10.1186/s12940-015-0051-6

Kim, S., Li, H., Jin, Y., Armad, J., Gu, H., Mani, S., & Cui, J. Y. (2023). Maternal PBDE exposure disrupts gut microbiome and promotes hepatic proinflammatory signaling in humanized PXR-transgenic mouse offspring over time. Toxicological Sciences: An Official Journal of the Society of Toxicology, 194(2), 209–225. 10.1093/toxsci/kfad056

Kim, Y.-M., Nam, I.-H., Murugesan, K., Schmidt, S., Crowley, D. E., & Chang, Y.-S. (2007). Biodegradation of diphenyl ether and transformation of selected brominated congeners by Sphingomonas sp. PH-07. Applied Microbiology and Biotechnology, 77(1), 187–194. 10.1007/s00253-007-1129-z

Knobloch, H. S., Charlet, A., Hoffmann, L. C., Eliava, M., Khrulev, S., Cetin, A. H., Osten, P., Schwarz, M. K., Seeburg, P. H., Stoop, R., & Grinevich, V. (2012). Evoked axonal oxytocin release in the central amygdala attenuates fear response. Neuron, 73(3), 553–566. 10.1016/j.neuron.2011.11.030

Kozlova, E. V., Chinthirla, B. D., Pérez, P. A., DiPatrizio, N. V., Argueta, D. A., Phillips, A. L., Stapleton, H. M., González, G. M., Krum, J. M., Carrillo, V., & Others. (2020). Maternal transfer of environmentally relevant polybrominated diphenyl ethers (PBDEs) produces a diabetic phenotype and disrupts glucoregulatory hormones and hepatic endocannabinoids in adult mouse female offspring. Scientific Reports, 10(1), 1–17. https://www.nature.com/articles/s41598-020-74853-9

Kozlova, E. V., Denys, M. E., Bishay, A. E., Luna, C. N., De Angelis, M., Campoy, L., Habbal, A., Lam, A. A., Luvsanravdan, N., Ghilenschi Colton, A., Carter, D., Müller, T., Schramm, K.-W., & Currás-Collazo, M. C. (2025). Maternal Thyroid Supplementation Prevents Autistic-relevant Social Behavior and Hypothalamic Oxytocin Depletion Produced by Developmental Exposure to Environmental Toxicants. In Neuroscience (No. biorxiv;2025.07.28.667293v1). bioRxiv. https://www.biorxiv.org/content/10.1101/2025.07.28.667293v1

Kozlova, E. V., Gonzalez, G. M., Denys, M. E., Bishay, A. E., Gutierrez, R., Reid, J., Krum, J. M., Lampel, G., Luvsanravdan, N., Rabbani, K. M., Tu, J., Campoy, L., Anchondo, L. M., Luna, C. N., Olomi, D. S., Monarrez, E., Carrillo, V., Tran, J. D., Platt, D., … Curras-Collazo, M. C. (2025). Enduring Autism-like Phenotypes and Deregulated Hypothalamic Prosocial Peptides After Early-Life Exposure to Indoor Flame Retardants in Male C57BL/6 Mice. In Neuroscience (No. biorxiv;2025.09.02.673882v1). bioRxiv. https://www.biorxiv.org/content/10.1101/2025.09.02.673882v1

Kozlova, E. V., Valdez, M. C., Denys, M. E., Bishay, A. E., Krum, J. M., Rabbani, K. M., Carrillo, V., Gonzalez, G. M., Lampel, G., Tran, J. D., Vazquez, B. M., Anchondo, L. M., Uddin, S. A., Huffman, N. M., Monarrez, E., Olomi, D. S., Chinthirla, B. D., Hartman, R. E., Kodavanti, P. R. S., … Curras-Collazo, M. C. (2022). Persistent autism-relevant behavioral phenotype and social neuropeptide alterations in female mice offspring induced by maternal transfer of PBDE congeners in the commercial mixture DE-71. Archives of Toxicology, 96(1), 335–365. 10.1007/s00204-021-03163-4

Larsson, M., Tirado, C., & Wiens, S. (2017). A meta-analysis of odor thresholds and odor identification in autism Spectrum Disorders. Frontiers in Psychology, 8, 679. 10.3389/fpsyg.2017.00679

Laue, H. E., Brennan, K. J. M., Gillet, V., Abdelouahab, N., Coull, B. A., Weisskopf, M. G., Burris, H. H., Zhang, W., Takser, L., & Baccarelli, A. A. (2019). Associations of prenatal exposure to polybrominated diphenyl ethers and polychlorinated biphenyls with long-term gut microbiome structure: a pilot study. Environmental Epidemiology (Philadelphia, Pa.), 3(1). 10.1097/EE9.0000000000000039

Li, C. Y., Dempsey, J. L., Wang, D., Lee, S., Weigel, K. M., Fei, Q., Bhatt, D. K., Prasad, B., Raftery, D., Gu, H., & Cui, J. Y. (2018). PBDEs altered gut microbiome and bile acid homeostasis in male C57BL/6 mice. Drug Metabolism and Disposition: The Biological Fate of Chemicals, 46(8), 1226– 1240. 10.1124/dmd.118.081547

Lin, Y. P., Thibodeaux, C. H., Peña, J. A., Ferry, G. D., & Versalovic, J. (2008). Probiotic Lactobacillus reuteri suppress proinflammatory cytokines via c-Jun. Inflammatory Bowel Diseases, 14(8), 1068– 1083. 10.1002/ibd.20448

Lord, C., Elsabbagh, M., Baird, G., & Veenstra-Vanderweele, J. (2018). Autism spectrum disorder. Lancet, 392(10146), 508–520. 10.1016/S0140-6736(18)31129-2

Mazzone, L., Dooling, S. W., Volpe, E., Uljarević, M., Waters, J. L., Sabatini, A., Arturi, L., Abate, R., Riccioni, A., Siracusano, M., Pereira, M., Engstrand, L., Cristofori, F., Adduce, D., Francavilla, R., Costa-Mattioli, M., & Hardan, A. Y. (2024). Precision microbial intervention improves social behavior but not autism severity: A pilot double-blind randomized placebo-controlled trial. Cell Host & Microbe, 32(1), 106–116.e6. 10.1016/j.chom.2023.11.021

Messer, A. (2010). Mini-review: polybrominated diphenyl ether (PBDE) flame retardants as potential autism risk factors. Physiology & Behavior, 100(3), 245–249. 10.1016/j.physbeh.2010.01.011

Minett, M. S., Quick, K., & Wood, J. N. (2011). Behavioral Measures of Pain Thresholds. Current Protocols in Mouse Biology, 1(3), 383–412. 10.1002/9780470942390.mo110116

Morais, L. H., Golubeva, A. V., Moloney, G. M., Moya-Pérez, A., Ventura-Silva, A. P., Arboleya, S., Bastiaanssen, T. F. S., O’Sullivan, O., Rea, K., Borre, Y., Scott, K. A., Patterson, E., Cherry, P., Stilling, R., Hoban, A. E., El Aidy, S., Sequeira, A. M., Beers, S., Moloney, R. D., … Cryan, J. F. (2020). Enduring Behavioral Effects Induced by Birth by Caesarean Section in the Mouse. Current Biology: CB, 30(19), 3761–3774.e6. 10.1016/j.cub.2020.07.044

Moy, S. S., Nadler, J. J., Perez, A., Barbaro, R. P., Johns, J. M., Magnuson, T. R., Piven, J., & Crawley, J. N. (2004). Sociability and preference for social novelty in five inbred strains: an approach to assess autistic-like behavior in mice. Genes, Brain, and Behavior, 3(5), 287–302. 10.1111/j.1601-1848.2004.00076.x

Mu, Q., Tavella, V. J., & Luo, X. M. (2018). Role of Lactobacillus reuteri in Human Health and Diseases. Frontiers in Microbiology, 9, 757. 10.3389/fmicb.2018.00757

Nomura, M., McKenna, E., Korach, K. S., Pfaff, D. W., & Ogawa, S. (2002). Estrogen receptor-beta regulates transcript levels for oxytocin and arginine vasopressin in the hypothalamic paraventricular nucleus of male mice. Brain Research. Molecular Brain Research, 109(1-2), 84–94. 10.1016/s0169-328x(02)00525-9

Oettl, L.-L., Ravi, N., Schneider, M., Scheller, M. F., Schneider, P., Mitre, M., da Silva Gouveia, M., Froemke, R. C., Chao, M. V., Young, W. S., Meyer-Lindenberg, A., Grinevich, V., Shusterman, R., & Kelsch, W. (2016). Oxytocin Enhances Social Recognition by Modulating Cortical Control of Early Olfactory Processing. Neuron, 90(3), 609–621. 10.1016/j.neuron.2016.03.033

Ongono, J. S., Dow, C., Gambaretti, J., Severi, G., Boutron-Ruault, M.-C., Bonnet, F., Fagherazzi, G., & Mancini, F. R. (2019). Dietary exposure to brominated flame retardants and risk of type 2 diabetes in the French E3N cohort. Environment International, 123, 54–60. 10.1016/j.envint.2018.11.040

Patisaul, H. B. (2017). Endocrine disruption of vasopressin systems and related behaviors. Frontiers in Endocrinology, 8, 134. 10.3389/fendo.2017.00134

Peng, Y., Ma, Y., Luo, Z., Jiang, Y., Xu, Z., & Yu, R. (2023). Lactobacillus reuteri in digestive system diseases: focus on clinical trials and mechanisms. Frontiers in Cellular and Infection Microbiology, 13, 1254198. 10.3389/fcimb.2023.1254198

Pfaffl, M. W. (2001). A new mathematical model for relative quantification in real-time RT–PCR. Nucleic Acids Research, 29(9), e45–e45. 10.1093/nar/29.9.e45

Poutahidis, T., Kearney, S. M., Levkovich, T., Qi, P., Varian, B. J., Lakritz, J. R., Ibrahim, Y. M., Chatzigiagkos, A., Alm, E. J., & Erdman, S. E. (2013). Microbial symbionts accelerate wound healing via the neuropeptide hormone oxytocin. PloS One, 8(10), e78898. 10.1371/journal.pone.0078898

Poutahidis, T., Kleinewietfeld, M., Smillie, C., Levkovich, T., Perrotta, A., Bhela, S., Varian, B. J., Ibrahim, Y. M., Lakritz, J. R., Kearney, S. M., Chatzigiagkos, A., Hafler, D. A., Alm, E. J., & Erdman, S. E. (2013). Microbial reprogramming inhibits Western diet-associated obesity. PloS One, 8(7), e68596. 10.1371/journal.pone.0068596

Qiu, H., Gao, H., Yu, F., Xiao, B., Li, X., Cai, B., Ge, L., Lu, Y., Wan, Z., Wang, Y., Xia, T., Wang, A., & Zhang, S. (2022). Perinatal exposure to low-level PBDE-47 programs gut microbiota, host metabolism and neurobehavior in adult rats: An integrated analysis. The Science of the Total Environment, 825, 154150. 10.1016/j.scitotenv.2022.154150

Raam, T., McAvoy, K. M., Besnard, A., Veenema, A. H., & Sahay, A. (2017). Hippocampal oxytocin receptors are necessary for discrimination of social stimuli. Nature Communications, 8(1), 2001. 10.1038/s41467-017-02173-0

Rigney, N., Campos-Lira, E., Kirchner, M. K., Wei, W., Belkasim, S., Beaumont, R., Singh, S., Suarez, S. G., Hartswick, D., Stern, J. E., de Vries, G. J., & Petrulis, A. (2024). A vasopressin circuit that modulates mouse social investigation and anxiety-like behavior in a sex-specific manner. Proceedings of the National Academy of Sciences of the United States of America, 121(20), e2319641121. 10.1073/pnas.2319641121

Robrock, K. R., Coelhan, M., Sedlak, D. L., & Alvarez-Cohent, L. (2009). Aerobic biotransformation of polybrominated diphenyl ethers (PBDEs) by bacterial isolates. Environmental Science & Technology, 43(15), 5705–5711. 10.1021/es900411k

Scattoni, M. L., Gandhy, S. U., Ricceri, L., & Crawley, J. N. (2008). Unusual repertoire of vocalizations in the BTBR T+tf/J mouse model of autism. PloS One, 3(8), e3067. 10.1371/journal.pone.0003067

Scheggia, D., Managò, F., Maltese, F., Bruni, S., Nigro, M., Dautan, D., Latuske, P., Contarini, G., Gomez-Gonzalo, M., Requie, L. M., Ferretti, V., Castellani, G., Mauro, D., Bonavia, A., Carmignoto, G., Yizhar, O., & Papaleo, F. (2020). Somatostatin interneurons in the prefrontal cortex control affective state discrimination in mice. Nature Neuroscience, 23(1), 47–60. 10.1038/s41593-019-0551-8

Scoville, D. K., Li, C. Y., Wang, D., Dempsey, J. L., Raftery, D., Mani, S., Gu, H., & Cui, J. Y. (2019). Polybrominated Diphenyl Ethers and Gut Microbiome Modulate Metabolic Syndrome–Related Aqueous Metabolites in Mice. Drug Metabolism and Disposition: The Biological Fate of Chemicals, 47(8), 928–940. 10.1124/dmd.119.086538

Sgritta, M., Dooling, S. W., Buffington, S. A., Momin, E. N., Francis, M. B., Britton, R. A., & Costa-Mattioli, M. (2019). Mechanisms Underlying Microbial-Mediated Changes in Social Behavior in Mouse Models of Autism Spectrum Disorder. Neuron, 101(2), 246–259.e6. 10.1016/j.neuron.2018.11.018

Shaw, K. A., Williams, S., Patrick, M. E., Valencia-Prado, M., Durkin, M. S., Howerton, E. M., Ladd-Acosta, C. M., Pas, E. T., Bakian, A. V., Bartholomew, P., Nieves-Muñoz, N., Sidwell, K., Alford, A., Bilder, D. A., DiRienzo, M., Fitzgerald, R. T., Furnier, S. M., Hudson, A. E., Pokoski, O. M., … Maenner, M. J. (2025). Prevalence and early identification of autism spectrum disorder among children aged 4 and 8 years - autism and Developmental Disabilities Monitoring Network, 16 sites, United States, 2022. MMWR Surveillance Summaries, 74(2), 1–22. 10.15585/mmwr.ss7402a1

Silverman, J. L., Yang, M., Lord, C., & Crawley, J. N. (2010). Behavioural phenotyping assays for mouse models of autism. Nature Reviews. Neuroscience, 11(7), 490–502. 10.1038/nrn2851

Takayanagi, Y., Yoshida, M., Bielsky, I. F., Ross, H. E., Kawamata, M., Onaka, T., Yanagisawa, T., Kimura, T., Matzuk, M. M., Young, L. J., & Nishimori, K. (2005). Pervasive social deficits, but normal parturition, in oxytocin receptor-deficient mice. Proceedings of the National Academy of Sciences of the United States of America, 102(44), 16096–16101. 10.1073/pnas.0505312102

Tamburini, S., Shen, N., Wu, H. C., & Clemente, J. C. (2016). The microbiome in early life: implications for health outcomes. Nature Medicine, 22(7), 713–722. 10.1038/nm.4142

Tan, Y., Singhal, S. M., Harden, S. W., Cahill, K. M., Nguyen, D.-T. M., Colon-Perez, L. M., Sahagian, T. J., Thinschmidt, J. S., de Kloet, A. D., Febo, M., Frazier, C. J., & Krause, E. G. (2019). Oxytocin receptors are expressed by glutamatergic prefrontal cortical neurons that selectively modulate social recognition. The Journal of Neuroscience: The Official Journal of the Society for Neuroscience, 39(17), 3249–3263. 10.1523/JNEUROSCI.2944-18.2019

Teffera, M., Veith, A. C., Ronnekleiv-Kelly, S., Bradfield, C. A., Nikodemova, M., Tussing-Humphreys, L., & Malecki, K. (2024). Diverse mechanisms by which chemical pollutant exposure alters gut microbiota metabolism and inflammation. Environment International, 190(108805), 108805. 10.1016/j.envint.2024.108805

Thirtamara Rajamani, K., Barbier, M., Lefevre, A., Niblo, K., Cordero, N., Netser, S., Grinevich, V., Wagner, S., & Harony-Nicolas, H. (2024). Oxytocin activity in the paraventricular and supramammillary nuclei of the hypothalamus is essential for social recognition memory in rats. Molecular Psychiatry, 29(2), 412–424. 10.1038/s41380-023-02336-0

Varian, B. J., Poutahidis, T., DiBenedictis, B. T., Levkovich, T., Ibrahim, Y., Didyk, E., Shikhman, L., Cheung, H. K., Hardas, A., Ricciardi, C. E., Kolandaivelu, K., Veenema, A. H., Alm, E. J., & Erdman, S. E. (2017). Microbial lysate upregulates host oxytocin. Brain, Behavior, and Immunity, 61, 36–49. 10.1016/j.bbi.2016.11.002

Varian, B. J., Poutahidis, T., Levkovich, T., Ibrahim, Y. M., Lakritz, J. R., Chatzigiagkos, A., Scherer-Hoock, A., Alm, E. J., & Erdman, S. E. (2014). Beneficial Bacteria Stimulate Youthful Thyroid Gland Activity. Journal of Obesity & Weight Loss Therapy, 4(2), 1–8. 10.4172/2165-7904.1000220

Vasudevan, N., Davidkova, G., Zhu, Y. S., Koibuchi, N., Chin, W. W., & Pfaff, D. (2001). Differential interaction of estrogen receptor and thyroid hormone receptor isoforms on the rat oxytocin receptor promoter leads to differences in transcriptional regulation. Neuroendocrinology, 74(5), 309–324. 10.1159/000054698

Vuong, H. E., & Hsiao, E. Y. (2017). Emerging roles for the gut microbiome in autism spectrum disorder. Biological Psychiatry, 81(5), 411–423. 10.1016/j.biopsych.2016.08.024

Wagner, S., & Harony-Nicolas, H. (2018). Oxytocin and Animal Models for Autism Spectrum Disorder. Current Topics in Behavioral Neurosciences, 35, 213–237. 10.1007/7854_2017_15

Wang, X., Hu, R., Lin, F., Yang, T., Lu, Y., Sun, Z., Li, T., & Chen, J. (2024). Lactobacillus reuteri or Lactobacillus rhamnosus GG intervention facilitates gut barrier function, decreases corticosterone and ameliorates social behavior in LPS-exposed offspring. Food Research International (Ottawa, Ont.), 197(Pt 1), 115212. 10.1016/j.foodres.2024.115212

Whitnall, M. H., Key, S., Ben-Barak, Y., Ozato, K., & Gainer, H. (1985). Immunocytochemical studies of the ontogeny of oxytocinergic and vasopressinergic neurons’. The Journal of Neuroscience: The Official Journal of the Society for Neuroscience.

Witchey, S. K., Al Samara, L., Horman, B. M., Stapleton, H. M., & Patisaul, H. B. (2020). Perinatal exposure to FireMaster® 550 (FM550), brominated or organophosphate flame retardants produces sex and compound specific effects on adult Wistar rat socioemotional behavior. Hormones and Behavior, 126, 104853. 10.1016/j.yhbeh.2020.104853

Woods, R., Vallero, R. O., Golub, M. S., Suarez, J. K., Ta, T. A., Yasui, D. H., Chi, L.-H., Kostyniak, P. J., Pessah, I. N., Berman, R. F., & LaSalle, J. M. (2012). Long-lived epigenetic interactions between perinatal PBDE exposure and Mecp2308 mutation. Human Molecular Genetics, 21(11), 2399–2411. 10.1093/hmg/dds046

Woolfe, G., & Macdonald, A. D. (1944). THE EVALUATION OF THE ANALGESIC ACTION OF PETHIDINE HYDROCHLORIDE (DEMEROL). The Journal of Pharmacology and Experimental Therapeutics, 80(3), 300–307. https://jpet.aspetjournals.org/content/80/3/300.short

Wu, W.-L., Wang, C.-H., Huang, E. Y.-K., & Chen, C.-C. (2009). Asic3(-/-) female mice with hearing deficit affects social development of pups. PloS One, 4(8), e6508. 10.1371/journal.pone.0006508

Yang, R., Ma, L., Peng, H., Zhai, Y., Zhou, G., Zhang, L., Zhuo, L., Wu, W., Guo, Y., Han, J., Jing, L., Zhou, X., Ma, X., & Li, Y. (2025). Microalgae-based bacteria for oral treatment of ASD through enhanced intestinal colonization and homeostasis. Theranostics, 15(6), 2139–2158. 10.7150/thno.103737

Zheng, J., Wittouck, S., Salvetti, E., Franz, C. M. A. P., Harris, H. M. B., Mattarelli, P., O’Toole, P. W., Pot, B., Vandamme, P., Walter, J., Watanabe, K., Wuyts, S., Felis, G. E., Gänzle, M. G., & Lebeer, S. (2020). A taxonomic note on the genus Lactobacillus: Description of 23 novel genera, emended description of the genus Lactobacillus Beijerinck 1901, and union of Lactobacillaceae and Leuconostocaceae. International Journal of Systematic and Evolutionary Microbiology, 70(4), 2782–2858. 10.1099/ijsem.0.004107

