## Supplementary Information for "Social and neuroendocrine phenotypes reprogrammed by endocrine-disrupting chemicals can be mitigated by *Limosilactobacillus reuteri* modulation of the gut microbiome–thyroid–oxytocin axis"

**Article Type**: Original paper

**Figure Number**: 7

**Word Count**: 10,515

### Declarations

#### Disclaimer

Research reported in this publication was supported by the National Institute Of Environmental Health Sciences of the National Institutes of Health under Award Numbers F31ES034304 and F31AI179030. The content is solely the responsibility of the authors and does not necessarily represent the official views of the National Institutes of Health.

**Funding**
This work was supported by a Danone North America Gut Microbiome, Yogurt and Probiotics Fellowship Grant and a University of California Office of the President UC-Hispanic Serving Institutions Doctoral Diversity Initiative, President's Pre-Professoriate Fellowship (UC-HSI DDI) and F31ES034304 (E.V.K.); a University of California Chancellor's Undergraduate Research Fellowship (M.E.D.); NIH/NIGMS R35GM124724 (A.H.); F31AI179030 (E.A.D.) and UCR Academic Senate grant (M.C.C.).

**Acknowledgements**

**Conflicts of interests/Competing interests**
The authors report no conflicts of interests and have no competing interests to declare.

**Ethics approval**
Care and treatment of animals was performed in accordance with guidelines from and approved by the University of California, Riverside Institutional Animal Care and Use Committee (AUP# 5, 20170026, 20200018 and 20210031).

**Consent to participate**
Not applicable.

**Consent for publication**
All authors reviewed and approved the final manuscript.

**Availability of Data and Material**
Not applicable.

**Code Availability**
Not applicable.

**Author Contributions**

**Conceptualization,** E.V.K., M.E.D., M.C.-C.; **Methodology**, E.V.K., M.E.D., E.D., R.L., V.P., A.H., M.C.-C.; **Validation**, E.V.K., M.E.D., A.E.B., E.D., R.L., C.N.L., A.L., V.P., A.H., M.C.-C.; **Formal Analysis**, E.V.K., M.E.D., A.E.B., E.D., R.L., C.N.L., A.L., V.P., A.H., M.C.-C.; **Investigation**, E.V.K., M.E.D., A.E.B., E.D., R.L., C.N.L., A.L., V.P., A.H., M.C.-C.; **Writing – Original Draft**, E.V.K., M.E.D., M.C.-C.; **Writing – Reviewing and Editing**, E.V.K., M.E.D., E.D., R.L., C.N.L., A.L., V.P., A.H., M.C.-C.; **Visualization**, E.V.K., R.L.; **Resources**, E.V.K., M.E.D., A.H., M.C.-C.; **Data Curation**, E.V.K., M.E.D., A.E.B., E.D., R.L., V.P., A.H., M.C.-C.; **Supervision**, E.V.K., A.H., M.C.-C.; **Project Administration**, E.V.K., M.E.D., A.H., M.C.-C.; **Funding Acquisition**, E.V.K., M.E.D., A.H., M.C.-C.

**Supplementary Table 1. Dam food and water intake, perinatal weight gain and litter parameters.**

| | VEH/CON | VEH/CON+ LR | DE-71 | DE-71+LR |
|---|---|---|---|---|
| **Maternal Parameters** | | | | |
| **n** | 4 | 4 | 4 | 4 |
| **Gestational food intake (GD19-PND 0)** | | | | |
| **Absolute (g/day)** | 8.0 ± 0.7 | 6.1 ± 0.9 | 6.4 ± 0.6 | 6.3 ± 0.3 |
| **Relative (g/day/pup)** | 1.4 ± 0.2 | 1.1 ± 0.1 | 1.1 ± 0.3 | 1.1 ± 0.1 |
| **Gestational weight gain (GD19-PND 0)** | | | | |
| **Absolute (g/day)** | 1.9 ± 0.4 | 2.3 ± 0.4 | 1.7 ± 0.1 | 2.2 ± 0.4 |
| **Relative (g/day/pup)** | 0.23 ± 0.02 | 0.18 ± 0.01 | 0.25 ± 0.04 | 0.18 ± 0.06 |
| **Postpartum weight gain (PND 1-10)** | | | | |
| **Absolute (g/day)** | 0.3 ± 0.09 | 0.2 ± 0.03 | 0.2 ± 0.05 | 0.3 ± 0.03 |
| **Relative (g/day/pup)** | 0.08 ± 0.02 | 0.05 ± 0.01 | 0.05 ± 0.01 | 0.06 ± 0.01 |
| **Litter Parameters** | | | | |
| ***n*** | 7 | 6 | 6 | 8 |
| **Litter size** | 6.1 ± 0.26 | 5.8 ± 0.48 | 6.3 ± 0.67 | 5.1 ± 0.72 |
| **Secondary sex ratio (M/F)** | 0.58 ± 0.08 | 0.45 ± 0.09 | 0.58 ± 0.03 | 0.46 ± 0.07 |

**Supplementary Figure 1**

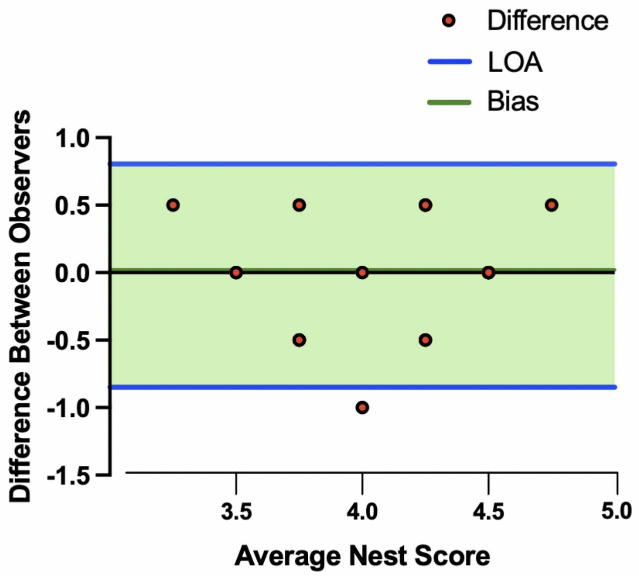


**Supplementary Figure 1. Bland-Altman plot for nest scores (Related to Figure 1).** Bland-Altman bias plot (mean±s.d.) was used to test the validity and reproducibility between two independent observers blind to exposure group. Analysis revealed a very small mean of the differences between observer scores (Bias, 0.02±0.42) and a precision measured as limits of agreement (LOA), average difference ± 1.96 standard deviation of the difference, of -0.80- 0.85, indicating negligible skewing by either observer.

**Supplementary Figure 2**

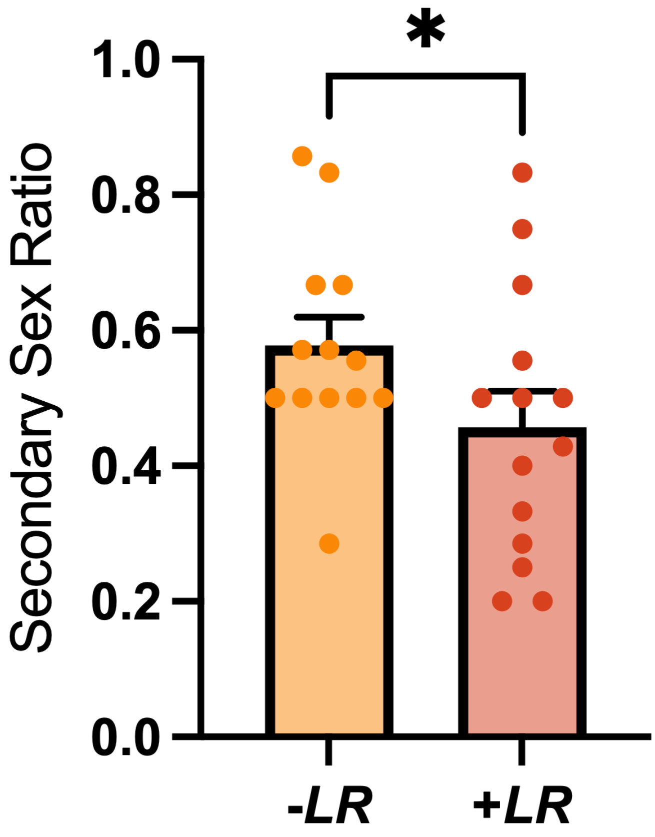


**Supplementary Figure 2. Secondary sex ratio (related to Supplementary Table 1).** Proportion of male over total offspring born to dams exposed to vehicle (CON) and 0.1 mg/kg DE-71 with or without LR supplementation. LR supplemented dams gave birth to a greater number of female pups as indicated by a lower secondary sex ratio. *indicates significantly different from -LR, *$p<.05$. *n*, 13-14 litters/group

**Supplementary Figure 3**

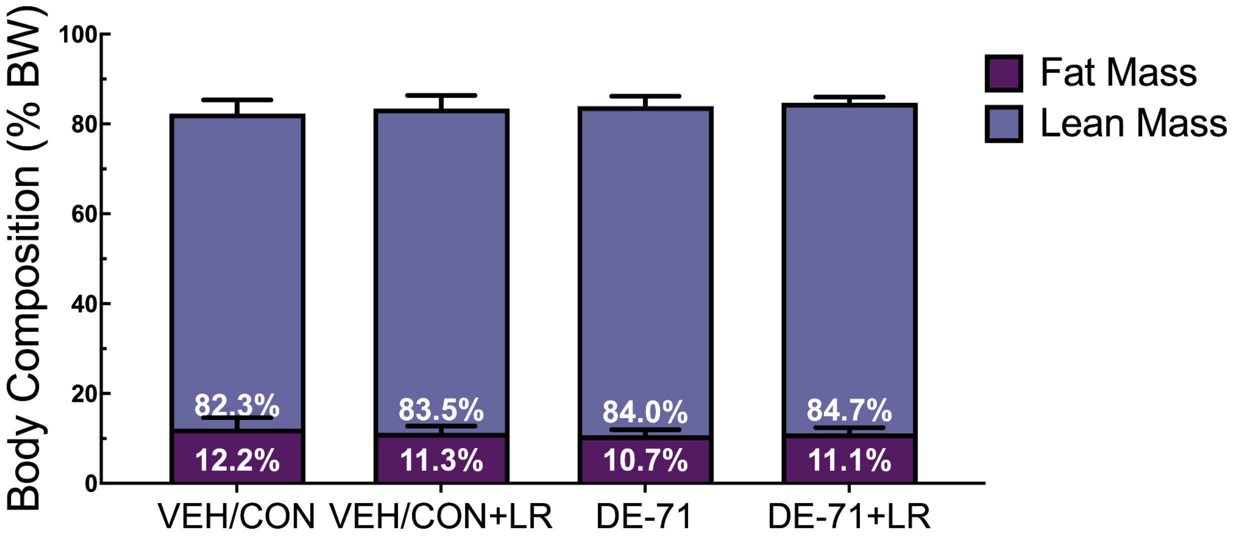


**Supplementary Figure 3 (Related to Figure 1). Dam body composition.** Fat mass and lean mass were measured at PND 23 using Echo MRI. Values are presented as a percentage of body weight. *n*, 6-8/group. Data represent mean ± SEM.

**Supplementary Figure 4**

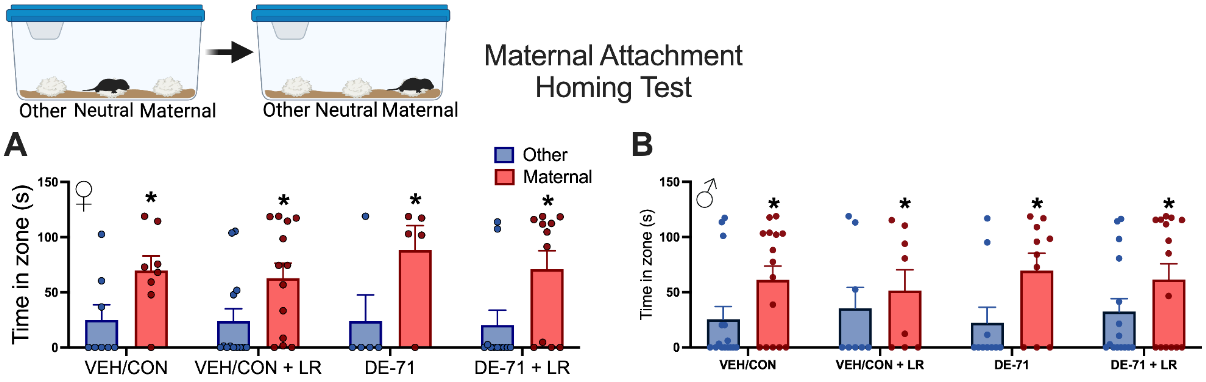


**Supplementary Figure 4 (related to Figure 1).** The maternal attachment and social recognition test was performed at PND 11. Pups were placed on neutral bedding and allowed to explore maternal or other dam bedding for 2 min. **(A)** Female offspring percent time in maternal zone and other zones. (**B**) Male offspring percent time spent in maternal and other zones. Values represent the mean of all same sex pups per litter. *n* females, 5-13 /group; n males, 8-15/group

**Supplementary Figure 5**

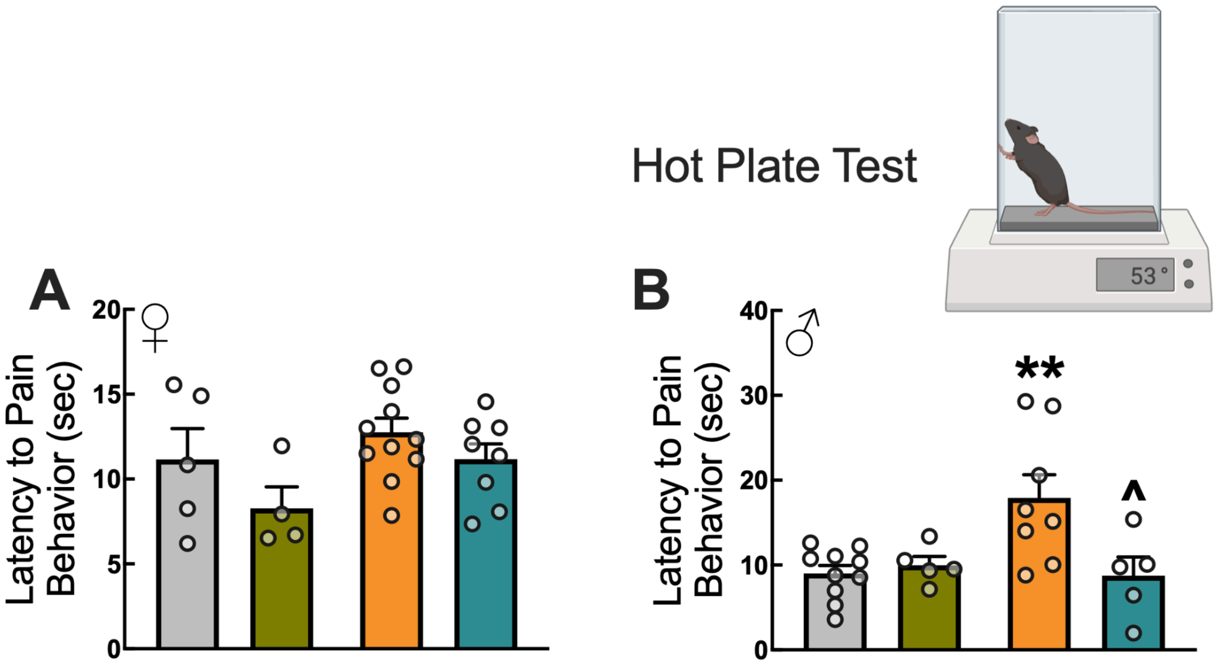


**Supplementary Figure 5.** Latency to withdraw on the hot plate test in (A) female offspring. (B) male offspring. *indicates significantly difference from VEH/CON, ***p*<.01; ^indicates significantly different from DE-71, ^*p*<.05. n, 4-11/group females, 5-10/group males

**Supplemental Figure 6**

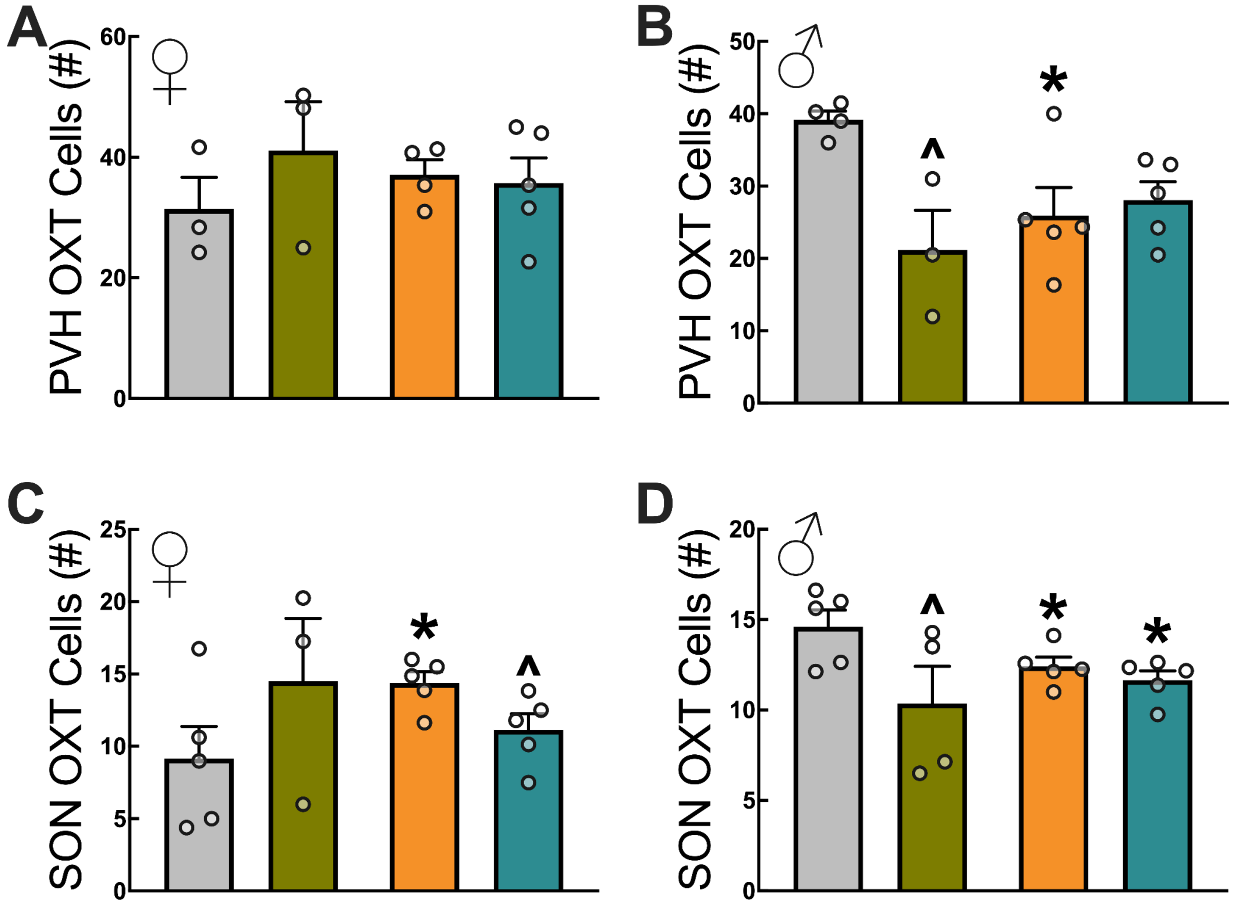


**Supplementary Figure 6. OXT cell counts in PVH and SON in male and female offspring (Related to Figures 4 and 5). (A)** Female PVH. **(B)** Male PVH. **(C)** Female SON. **(D)** Male SON. *indicates significantly difference from VEH/CON, *$p$<.05; ^indicates significantly different from corresponding unsupplemented group, ^$p$<.05. $n$, 3-5/group females, 4-5/group males

**Supplementary Figure 7**

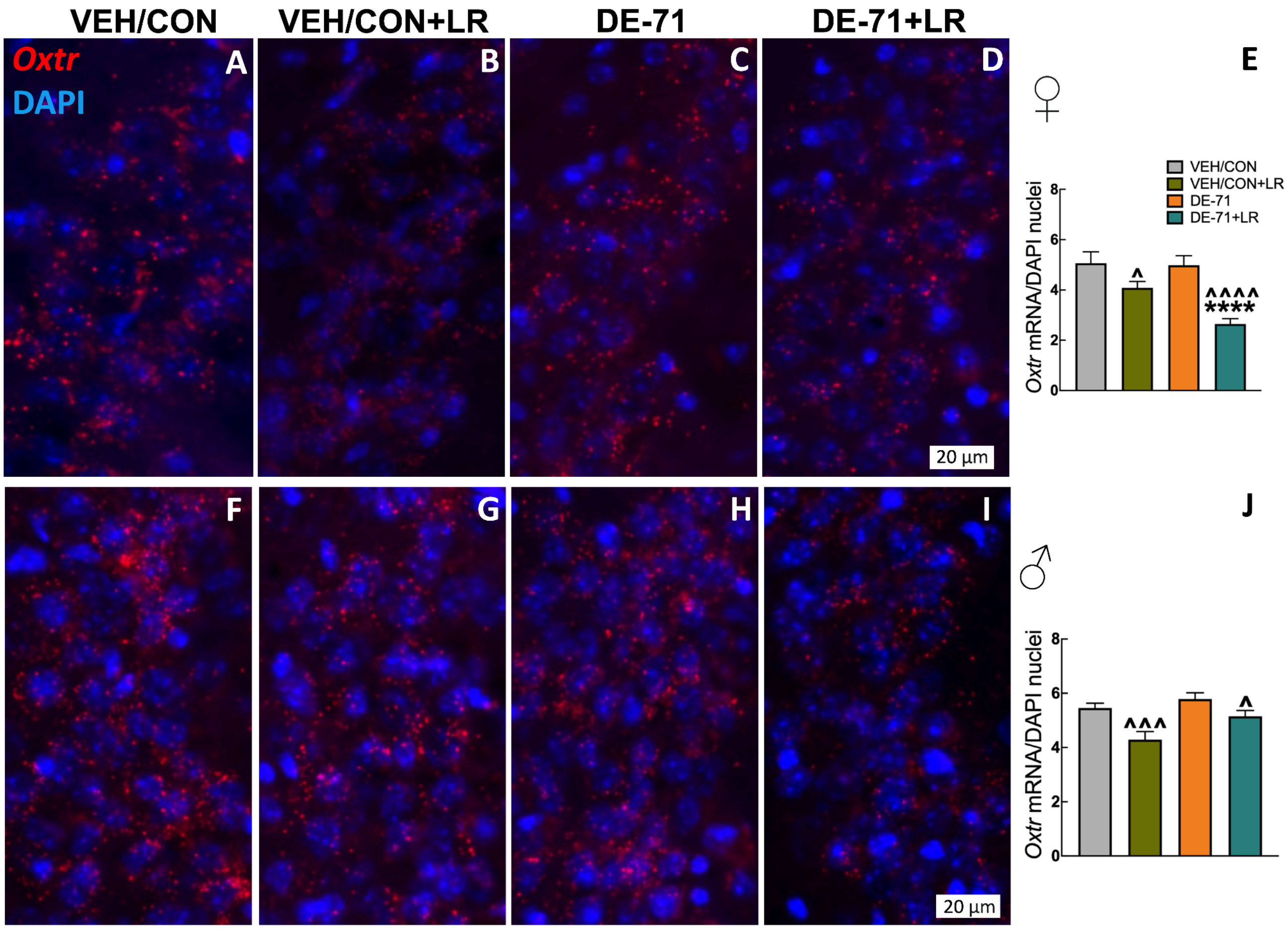


**Supplementary Figure 7. LR downregulates *Oxtr* in the dorsal anterior CA2/CA3 subfield of the hippocampal of both sexes.** Representative micrographs of RNA *in situ* hybridization, of *Oxtr* on DAPI-positive cells in females (**A-D**) and males (**F-I**). Mean group values of *Oxtr* mRNA transcript counts on DAPI+ cells in female (**E**) and male CA2 (**J**). *statistical difference vs VEH/CON, *$p$<0.05, ***$p$<0.001, ****$p$<0.0001. ^statistical difference vs DE-71, ^$p$<0.05, ^^^^$p$<0.0001. $n$, 3-5/group; 4-5/group. Scale bar, 20 µm.

**Supplementary Figure 8**

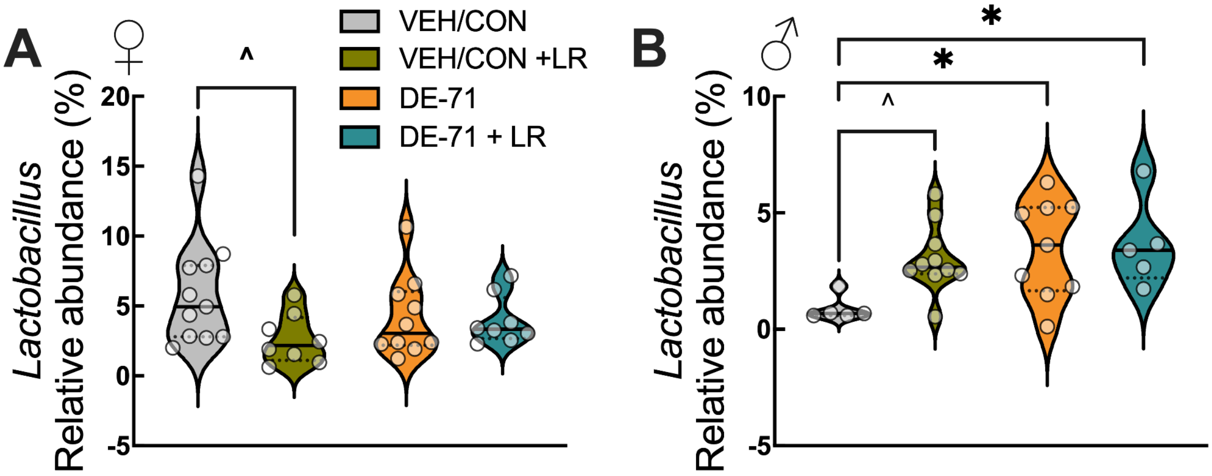


**Supplementary Figure 8. Relative abundance of *Lactobacillus* in Cohort 2 offspring in adulthood(Related to Figure 6). (A)** Relative abundance of *Lactobacillus* in female offspring. (**B**) Relative abundance of *Lactobacillus* in male offspring. *statistical difference vs VEH/CON, *$p$<0.05; Females $n$, 8-11/group; males $n$, 5-10/group
